# Early Electrophysiological Aberrations in the Hippocampus of the TgF344-AD Rat Model as a Potential Biomarker for Alzheimer’s Disease Prognosis

**DOI:** 10.1101/2022.07.01.498373

**Authors:** Faraz Moradi, Monica van den Berg, Morteza Mirjebreili, Lauren Kosten, Marleen Verhoye, Mahmood Amiri, Georgios A. Keliris

## Abstract

The hippocampus is thought to guide navigation and has an essential contribution to learning and memory. Hippocampus is one of the brain regions impaired in Alzheimer’s disease (AD), a neurodegenerative disease with progressive memory impairments and cognitive decline. Although successful treatments for AD are still not available, developing new strategies to detect AD at early stages before clinical manifestation is crucial for timely interventions. Here, we investigated in the TgF344-AD rat model the classification of AD-transgenic rats versus Wild-type littermates (WT) from electrophysiological activity recorded in the hippocampus of freely moving subjects at an early, pre-symptomatic stage of the disease (6 months old). To this end, recorded signals were filtered in two separate frequency regimes namely low frequency LFP signals and high frequency spiking activity and passed to machine learning (ML) classifiers to identify the genotype of the rats (TG vs. WT). For the low frequency analysis, we first filtered the signals and extracted the power spectra in different frequency bands known to carry differential information in the hippocampus (delta, theta, slow- and fast-gamma) while for the high frequency analysis, we extracted spike-trains of neurons and calculated different distance metrics between them, including Van Rossum (VR), Inter Spike Interval (ISI), and Event Synchronization (ES). These measures were then used as features for classification with different ML classifiers. We found that both low and high frequency signals were able to classify the rat genotype with a high accuracy with specific signals such as the gamma band power, providing an important fraction of information. In addition, when we combined information from both low and high frequency the classification was boosted indicating that independent information is present across the two bands. The results of this study offer a better insight into how different regions of the hippocampus are affected in earlier stages of AD.

## Introduction

Alzheimer’s disease (AD) is a neurodegenerative disorder affecting millions of people each year and has become one of the biggest socioeconomic burdens to societies. In the United States alone, over 5.7 million Americans were living with AD in 2018, and estimates indicate that this amount will increase to nearly 14 million by 2050^1^. The hallmarks of AD include the excessive aggregation of amyloid-β (Aβ) and tau proteins in brain areas including the hippocampus and adjacent mediotemporal cortex leading to progressive cognitive decline and, ultimately, dementia^2^. One of the first symptoms in AD is spatial memory impairment and wandering behavior with these symptoms occurring in over 60% of the patients^3^. Regardless of the many efforts and significant progress scientists and physicians have made in understanding AD, its leading cause and prognosis remain uncertain^2^. Current treatment methods can only decelerate AD progression especially if used in relatively early stages of AD. However, cognitive symptoms that are the leading trigger for AD diagnosis appear only at later stages of the disease timeline. Therefore, it has become essential to develop new strategies for the detection and prognosis of AD at early stages before the clinical manifestation of the disease^4^.

Hippocampus is a vital brain area that is involved in numerous cognitive functions including spatial navigation, learning, and memory^5^. Over several decades, investigations of hippocampal function have focused heavily on neurophysiological network activity, including the gamma and theta rhythms^6^. It has been demonstrated that theta oscillations, ranging in frequency from 4 to 12 Hz, are linked to a variety of cognitive processes and have an important contribution to learning and memory as observed in a wide range of experiments in both humans and animals^6–9^. Gamma oscillations, on the other hand, are characterized by activity in the frequencies ranging from 30 to 100 Hz^10^ and have been linked to functions such as sensory binding^11^, selective attention^12–14^, transient neuronal assembly formation^15^, and information transmission and storage^16–18^. Moreover, both types of oscillations, that are part of the low-frequency component of electrophysiological recordings called local field potentials (LFP; ∼ 0 - 150 Hz), have been identified as critical components in the consolidation of memories and the performance of executive functions^6,19– 24^. Also, another type of oscillation is high-voltage spindles (HVSs). They range from 5 to 13 Hz and have a characteristic spike-and-wave shape which has been studied mainly in somatosensory systems^25,26^, and only a few recent studies have seriously investigated their occurrence in the basal ganglia (BG)^27,28^. Theta and gamma oscillations do not exist in isolation; instead, they interact and co-occur in several areas of the brain^29^. A specific type of cross-frequency coupling (CFC), named phase-amplitude coupling (PAC), is characterized by the coupling between the phase of the theta and the amplitude of the gamma rhythms^26^. Moreover, research indicates that PAC in the hippocampus may play an essential role in learning and memory^29^. For humans, the magnitude of the coupling is found to be positively associated with cognitive processes^29–31^. Furthermore, PAC has been proposed to play a key functional role in learning task performance^32^, facilitation of inter-area communication^33,34^ and can serve as a memory buffer^31^. In relation to AD, there have been numerous articles suggesting that the oscillatory activity including power and/or coupling in theta^35–37^ and gamma^35,38^ rhythms is altered in AD patients, and consistent changes were also found in animal experiments using AD models^39,40^.

Beyond disturbances in the theta and gamma rhythms in the hippocampus and medial entorhinal cortex (MEC), many research papers also identified AD related disruptions in the activity of single neurons such as place cells and grid cells^39–41^. Place cells are neurons that selectively respond to specific location in the environment (Place field) first discovered by O’Keefe in CA1 region^52^. The activity of single neurons is characterized by discrete events called action potentials or spikes which are considered as the fundamental element of neuronal communication and can be extracted from the high-frequency component of electrophysiological recordings (> 300 Hz). One or more spikes in a sequence are generated whenever a single neuron is stimulated or even at rest (spontaneous activity), and a sequence of spikes forms what is referred to as a spike train^42^. It is generally accepted that the spike responses of each neuron are governed from two main information processing paradigms: rate coding and/or temporal coding^43^. Measuring the synchrony of two sets of spike trains has been demonstrated to carry a lot of information in several different contexts. For example, spike train distance metrics were used to estimate the reliability of responses across repeated presentations of the same stimulus and to evaluate the information flow among coupled neurons^44,45^. Furthermore, some recent studies reported the application of measuring spike train distance metrics for the classification of tactile afferents using artificial spike sequences^46^, raising the possibility that such metrics could potentially be used for the classification of AD in animal models.

With the rise of machine learning (ML) and deep learning (DL), numerous applications have been implemented towards the detection and classification of brain diseases. Many models used high dimensional features from various medical imaging techniques such as magnetic resonance imaging (MRI) and positron emission tomography (PET) for the prediction and diagnosis of AD. To this end, they employed classical ML methods like support vector machine (SVM) and k-nearest neighbor (k-NN), but also DL models such as convolutional neural networks (CNN) and recurrent neural networks (RNN). For example, Lee *et al*. ^47^ has proposed a new model to predict AD in which RNN was used considering multimodal features extracted from MRI and longitudinal cerebrospinal fluid (CSF) data. El-Sappagh et al. ^48^ used ensemble ML classifiers based on Random Forest (RF) to develop an AD diagnosis and progression detection model on 11 MRI modalities. Venugopalan et al. ^49^ used different models of ML (SVM, RF, and k-NN) and DL models using a fusion of multiple data modalities like MRI and genetic single nucleotide polymorphisms (SNPs) to classify patients into AD, mild cognitive impairment (MCI), and control groups. However, even though the above-mentioned methods demonstrated considerable success, in their great majority, they do not provide insights regarding the underlying disturbances at the local neuronal network and single neuron level.

In this study, we used a very promising rat model of AD that demonstrates all the pathophysiological hallmarks present in AD patients (Tg-F344-AD) and compared it with littermate wild-type (WT) controls. We put forward the hypothesis that local neuronal network features measured with electrophysiological recordings in the hippocampus of freely moving rats at an early time-point during disease progression, when amyloid plague accumulation is only at the onset (6 months old), can be used in combination ML-based methods to classify the rats into the two respective groups, namely AD and WT. Further, we were interested to identify which of these signals better classify the rats and if classification with a combination of signals can be advantageous. Specifically, we considered both low-frequency LFP signals as well as high-frequency spiking activity. To see the effect of low-frequency information in classification, we performed time-frequency analysis with focus on four specific LFP frequency bands (delta, theta, slow gamma, fast gamma) as well as CFC measures such as PAC. On the other hand, for the high-frequency analysis, we first used spike sorting algorithms to extract the spike trains of neurons and then computed different spike-train distance metrics such as Van Rossum (VR), Inter Spike Interval (ISI), and Event Synchronization (ES). We then used those measures as classification features to investigate the performance of different ML-classifiers across the two groups. We found that specific low and high frequency signals used with chosen ML methods could provide classification accuracies over 80% and that combining information from both could boost the accuracy to 100%. Our findings provide evidence that the disruption in local neuronal signaling in the hippocampus of an AD rat model could be used at a very early time-point of disease progression to classify between AD and WT rats and thus provide promise for the development of novel disease biomarkers for AD.

## MATERIALS AND METHODS

### Animals and ethical statement

All procedures were in accordance with the guidelines approved by the European Ethics Committee (decree 2010/63/EU) and were approved by the Committee on Animal Care and Use at the University of Antwerp, Belgium (approval number: 2019-06). Electrophysiological experiments were performed in 6-month-old TgF344-AD rats (N=5) and wild-type littermates (N=4). Rats were group-housed prior to head-stage implantation but housed separately afterward. All animals were kept on a reversed, 12h light/dark cycle, with controlled temperature (20 – 24°C) and humidity (40-60%) conditions. Standard food was provided ad libitum. Animals were water-deprived 24h prior to the habituation and acquisition of the LFP data. After the animal performed the task in the linear track, water was provided for 30 minutes.

### Chronic hippocampal electrophysiological measurements

#### Surgical procedure

For the implantation of the electrodes, animals were anesthetized using isoflurane (at 1.0 L/min, induction 5% and maintenance 2-3% isoflurane). Animals were placed in a stereotaxic frame and a craniotomy was made above the right dorsal hippocampus (AP -3.00, ML 2.50). A 16-channel laminar electrode (E16+R-100-S1-L6 NT, Atlas Neuro-engineering, Belgium) with internal reference was placed into the dorsal hippocampus by penetrating the dura (Fig. 1A). The depth of the recording sites was identified by the layer-specific local field potentials (LFP) of the hippocampus (DV 2.5-3.5 mm). The craniotomy was sealed with a sterile silicone gel (Kwik-Cast, WPI). Stainless steel screws were drilled into the skull overlaying the olfactory bulb, left hippocampus and cerebellum, of which the latter served as a ground electrode. Two EMG wires were stitched in the neck muscle in order to record EMG activity. The implant was covered in several layers of dental cement (Stoelting, Co, Dublin, Productnumber 50000,) and the wound was closed. Rats were allowed to recover for at least 7 days.

**Fig 1.**
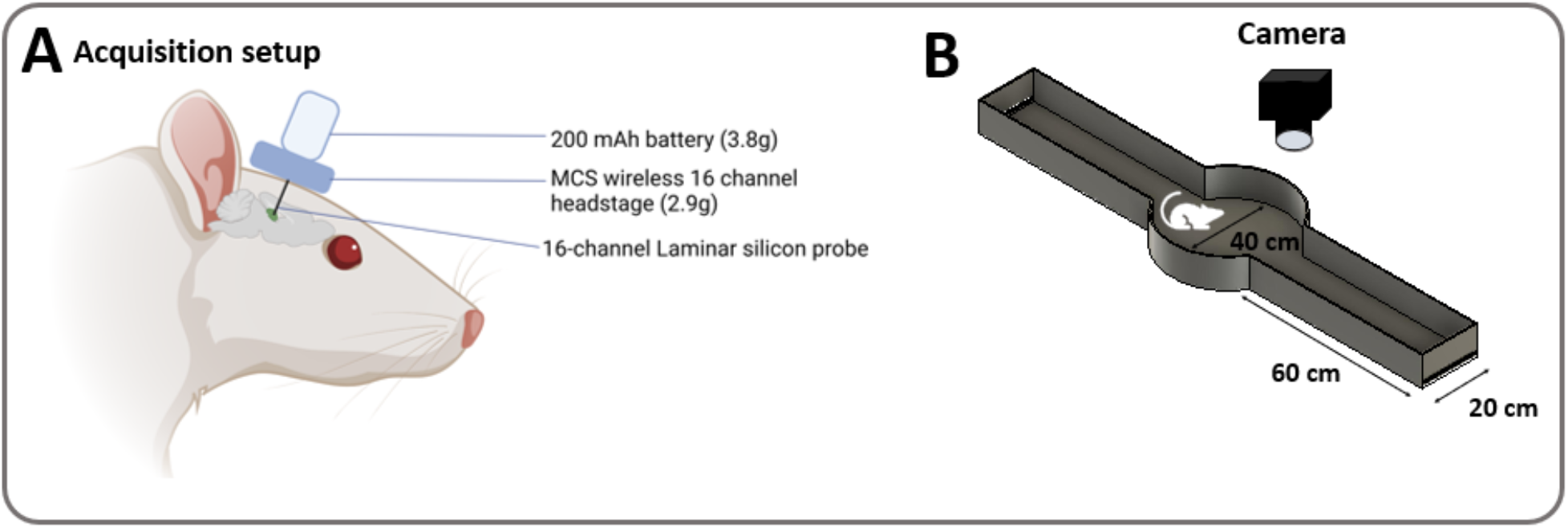
A. Graphical representation of the wireless acquisition system used to record LFP’s and neuronal activity. A head-stage was attached to the laminar silicon probe which was placed in the right dorsal hippocampus. The head-stage converts the analog signals recorded from the brain to digital signals, which were then send to the receiver hardware. The head-stage was powered by a small battery mounted on top of the head-stage to allow maximum freedom of movement. B. Graphical representation of the track where the animals freely moved during the recordings. It consisted of a circular area in the middle with a diameter of 40 cm and two rectangular arms of 20×60 cm. At the end of each arm, rewards were periodically provided to motivate the animals to move back and forth.

#### Linear track acquisition

Animals were habituated to the behavioral room and linear track (Fig. 1 B) for 1 day, after which the rats were trained for 5 consecutive days to walk along the linear track. Cups with sugar water at the end of the arms were used to motivate the animals to walk back and forth. Habituation sessions and training sessions lasted 20-30 min. The combined electrophysiological and behavioral acquisition was performed during two, or three consecutive days, based on the performance (active time) of the rats in the linear track. A wireless head stage (W2100, MultiChannel Systems, Germany) was connected to the laminar electrode, 15 to 30 minutes prior to start of the experiment in the linear track. After the habituation to the acquisition setup, animals were placed in the center of the track and the electrophysiological recordings were started. The movements of the animal were recorded using a camera mounted on the ceiling above the linear track, and the freely available Bonsai acquisition software (Bonsai — Open Ephys (open-ephys.org)) was used to detect the animal position over time. An acquisition session ended when the animal was not moving around for more than 5 minutes.

### Data Analysis

#### Preprocessing of LFP signals

Data was recorded from the 2 EMG channels and the laminar electrode consisting of 14 intracranial channels. The top three intracranial electrodes were located outside the hippocampus and were not used in our analysis. The remaining 11 intracranial channels were used for LFP analysis and spike sorting. To prevent the effect of impedance differences between electrodes on LFP, the LFP power for each electrode was normalized to the power of the 0.5-1 Hz band. The first and last 60 sec of each recording session were excluded to eliminate possible artifacts on the neural activity patterns associated with handling or moving into or out of the linear track. The LFP signals were extracted from the broad-band signal (sampling rate = 10 kHz) using a 3rd-order Butterworth filter (MATLAB function *filtfilt*) with a low cut-off frequency of 0.1 Hz and a high cut-off frequency of 300 Hz. After filtering the signal was downsampled to 1000 Hz and stored for further analysis. Out of 20 recording sessions from 9 rats (n=4 WT rat, n=5 TG rat), two of the samples (WT3_D1 and TG1_D2) were excluded from our further analysis because of excessive noise.

#### Power spectrum analysis

The power spectrum density (PSD) was estimated using a 2-second window size (MATLAB function *pwelch*). We computed the power between frequencies 1–100 Hz and normalized it to the power between 0.5-1 Hz as also indicated above. Then, the power at four individual frequency bands namely delta (1-4 Hz), theta (4-12 Hz), slow gamma (30-50 Hz), and fast gamma (50-100 Hz) bands was used for further analysis^40^.

#### Time-frequency analysis of power and frequency coupling

The Chronux^50^ toolbox in MATLAB (function *mtspecgramc*) was used for the time-frequency analysis. As the duration of recordings for each sample was between 17 to 30 minutes, we have chosen a snippet of 15 minutes (900 seconds) from each recording. The toolbox has parameters like window and step size, tapers which were tuned to obtain proper representations (window and step size = 15, taper = [5 9]). For computing cross-frequency coupling between desired phase and amplitude ranges, the Buzcode toolbox^51^ from Buzsaki Lab was used. The modulation index of phase-amplitude between desired ranges was computed for all data from each rat. Each representation is the result of averaging across all other channels.

#### Detection of sleep spindles

In our analysis, 15 min of recordings were used for all analyses. Automatic detection of high-voltage spindles (HVSs) was identified in the hippocampus when rhythmic negative deflections lower than −0.3 mV occurred while instantaneous power between 6 Hz and 12 Hz was more than double instantaneous delta power for 2 s or longer. The average time-frequency spectrum was calculated within frequencies between 0-20Hz for each animal.

### Classification Models

The rat’s hippocampus structure is depicted in Fig. 2A. This brain region receives information from various parts of the brain and is believed to support information about the spatial representation of our environment^52^. The most cortical input comes from the entorhinal cortex (EC)^53^. Encoded information transfers from EC → to CA1 DG via the perforant path (synapse 1), DG → CA3 via mossy fibres (synapse 2), CA3 → CA1 via schaffer collaterals (synapse 3) which together they form the trisynaptic circuit^54^. Therefore, we have decided to do the recording from this brain region as an essential organ in navigation and memory.

**Fig. 2.**
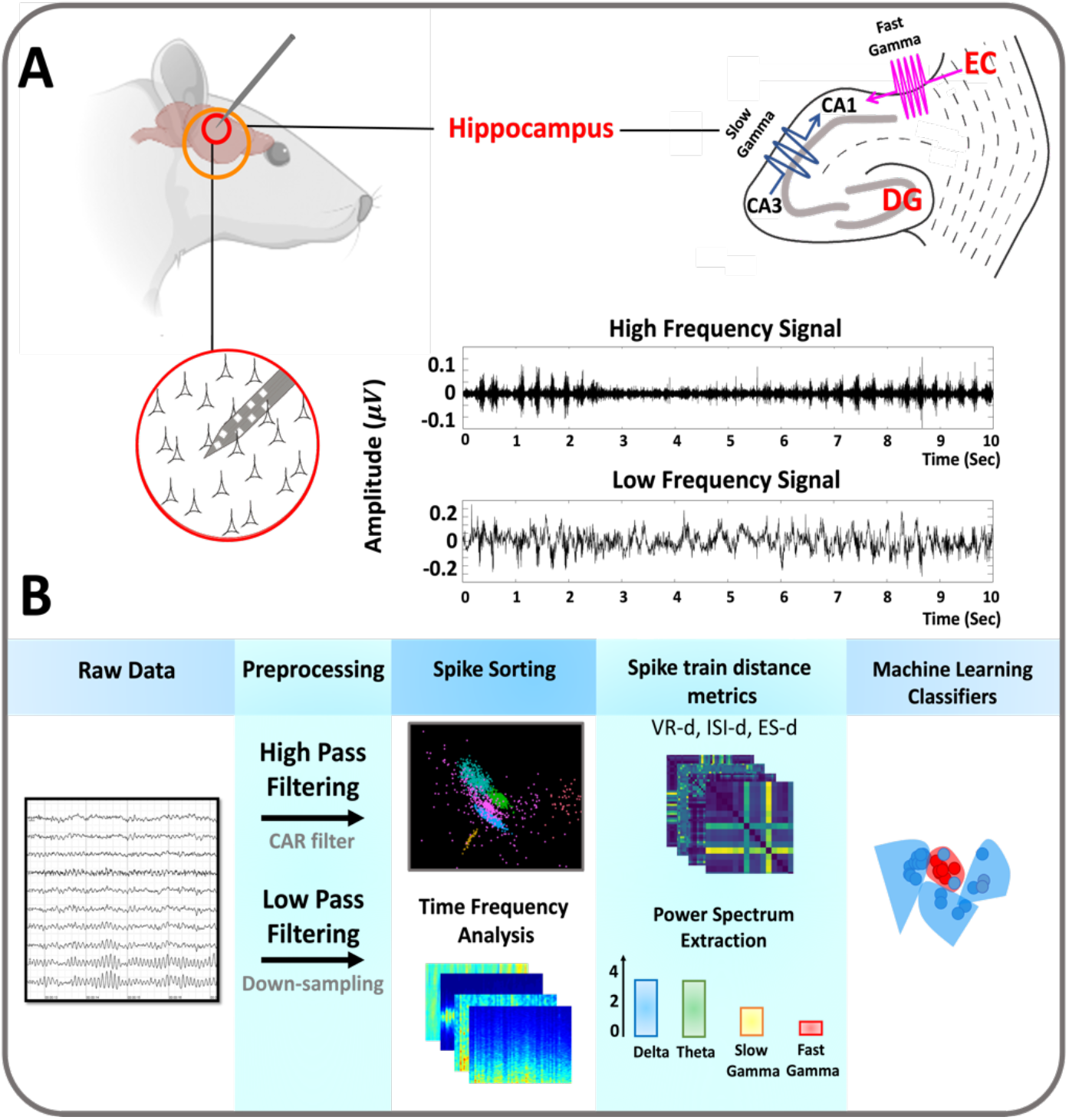
Representation of the recording and ML models. **A)** The relative positions of Hippocampus, entorhinal cortex (EC) and dentate gyrus (DG) formation are illustrated. Fast and slow gamma oscillations reflect inputs from medial EC and CA3, respectively. Graphs show representative data recorded in the hippocampus of one rat. The spike sorting algorithm runs on the high-frequency signal (band-pass filtered between 300 and 4900 Hz) of the raw data and uses common average reference (CAR) filter to remove mutual noise among channels. The low-frequency signal (band-pass filtered between 0.1 and 300 Hz) of the raw data was analyzed using time-frequency methods to extract desired frequency bands and then perform the classification task. **B)** Graphical representation of the two ML models. In the high-frequency model, spike sorting was performed after data acquisition and preprocessing to obtain the units spike train responses. Next, spike train distance and synchrony metrics were calculated, PCA was applied for dimension reduction and finally this information was fed into the ML classifiers that distinguished TG and WT rats. In the low-frequency model, the primary step was to analyze the time-frequency plots and then estimate the average power for individual frequency bands. By obtaining the most compelling features, the classification of TG and WT rats was then accomplished.

In Fig. 2B, the steps undertaken for the classification models are shown. We have put forward two different pipelines for the classification task: 1. Using Low-Frequency Signals (LFS) and 2. Using High-Frequency Signals (HFS). Moreover, we also combined the information from both these models to estimate the total performance of using these local network features for classification. In the LFS model, the average power spectrum density (PSD) is computed for the frequency bands of interest (delta, theta, slow gamma, fast gamma) and then the feature importance method was used by calculating the Mean Decrease in Impurity (MDI) to determine the effectiveness of each frequency band in the classification of TG and WT rats. The LFP data from each rat comprises the model input and is defined as *X* (*X* ∈ ℝ^*timepoint*×*Channel*^). The HFS model comprises of three main steps: (1) Running spike sorting algorithms using Tridesclous sorter^55^ and SpikeInterface^56^ toolboxes to isolate spike responses of neural activities; (2) Extracting spike train distance and synchrony metrics including Van Rossum (VR-d), Inter Spike Interval (ISI-d), and Event Synchronization (ES-d) for each TG and WT group; and (3) Applying principal component analysis (PCA) to reduce the dimensionality of the feature space.

### 1. Classification Using Low-Frequency Signals (LFS)

#### Spectral Power Model

Further analysis on the LFP signal was performed to investigate how TG and WT can be distinguished in low-frequency features using each frequency band’s mean power. In this case, the average power of delta (1-4 Hz), theta (4-12 Hz), slow gamma (30-50 Hz), and fast gamma (50-100 Hz) are extracted for every rat as shown in Fig. 3. Therefore, the shape of the extracted feature matrix is *S*_*WT*_ ×4 and *S*_*TG*_ ×4 where *S*_*WT*_ (*S*_*TG*_) is the number of samples for the WT (TG) group. Also, we have analyzed a scenario in which each frequency band is removed one by one from the input features of the ML classifier. Consequently, the shape of each feature matrix is decreased to *S*_*WT*_ ×1 and *S*_*TG*_ ×1 in the final step. This removal of each frequency band is used to demonstrate how different hippocampal neural oscillations affect the model’s performance. Due to the small number of samples, it was not possible to use the commonly used approach that shuffles the data from all samples and then divides them into training, validation, and test sets. Instead, we have used leave-one-subject-out cross-validation (LOSO-CV). In this approach, the number of folds is equal to the number of recording sessions. Thus, the ML classifier is applied once for each sample using all other samples as a training set and the selected sample as a single-item test set. Therefore, we have separated each sample’s spike train manually as a test set, and the training was performed using all the remaining spike train features. This process was continued until every recording session was used as a test set.

**Fig. 3.**
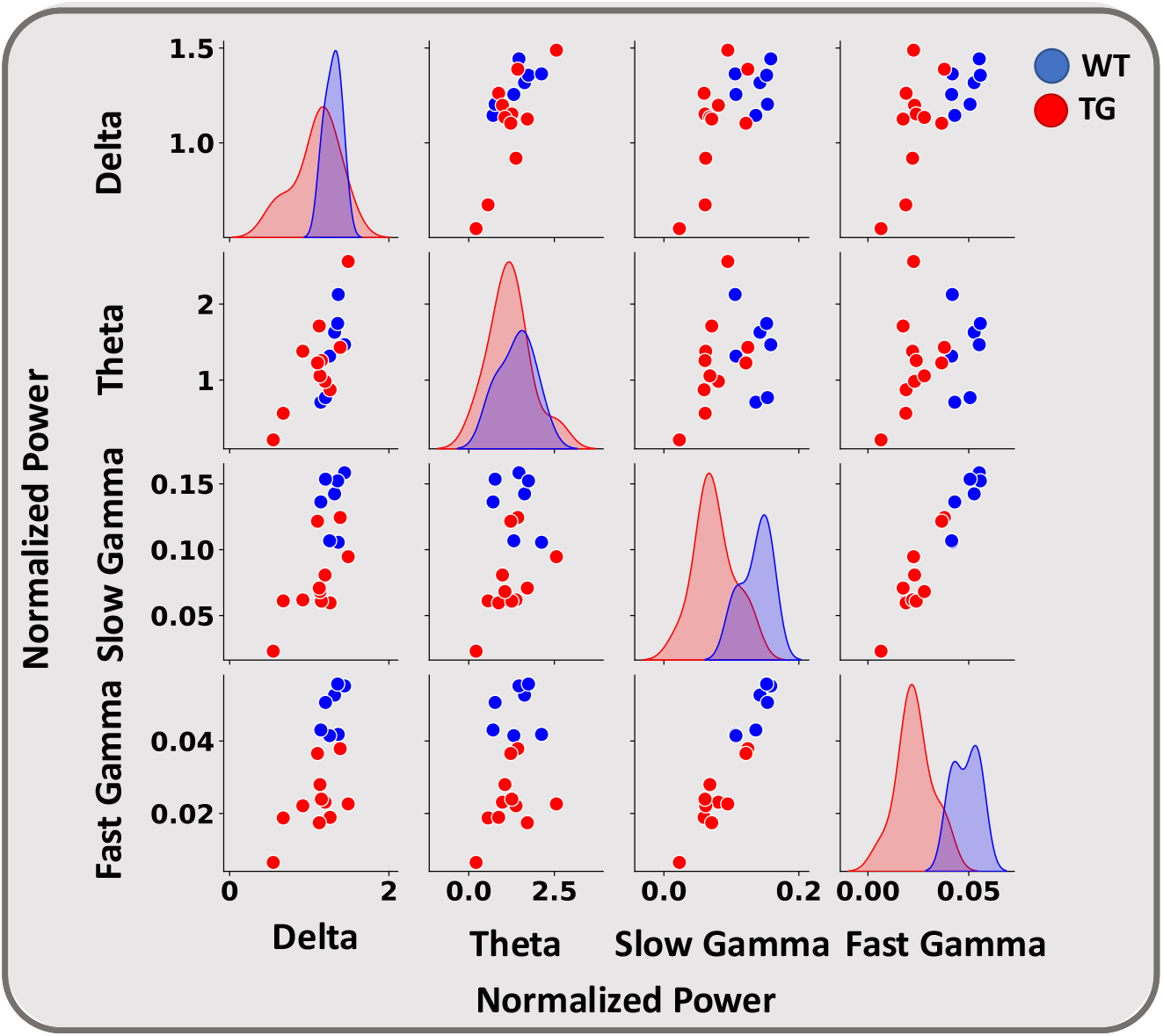
Pairwise comparison of the normalized power in delta (1-4 Hz), theta (4-12 Hz), slow gamma (30-50 Hz), and fast gamma (50-100 Hz) for TgF344-AD (TG) and Wildtype (WT) groups. Each point represents the average PSD of a single sample in the specified band. The diagonal plots are the univariate distributions that show the amount of overlap between the distribution of two groups in individual frequency bands. The other plots compare WT and TG for specified frequency bands. Blue/Red points are recording session (samples) of WT (n = 7) /TG (n = 11), respectively.

#### Feature Importance

The feature importance gives an abstract view about the role of each feature. Although this study aims to build a classifier to discriminate between TG and WT, it is also essential to quantify the importance of each input feature which in turn shows those frequency bands that have been changed in the AD rats. For this purpose, a random forest (RF) classifier was trained with all four frequency bands, and Mean Decrease in Impurity (MDI) has been used as a measure for evaluating the importance of each feature. The MDI calculates each feature importance as the sum over the number of splits (across all trees) that include the feature, proportionally to the number of samples it splits.

### 2. Classification Using High-Frequency Signals (HFS)

#### Spike Sorting and Multi-unit Extraction

Pre-processing for spike sorting included high-pass filtering with a cutoff at 300Hz and then low-pass filtering with a cutoff at 4900Hz. Also, common reference removal (CAR) filter was used to increase the signal-to-noise ratio.

Then, we used the python-based package Tridesclous Sorter, an automatic spike-sorting toolbox that groups spike responses into separate units based on their similarity in shape and features. We have called these spiking activities “multi-unit,” as neuron isolation from single electrodes on a laminar probe is difficult and thus these may contain spike responses from a few neighboring neurons. Another python-based package, and namely the Spike Interface toolbox has been used for manual inspection and curation of multi-units. We have defined three operators for each of the steps: Band-pass filter (⊛), Common average reference filter (⊕), Tridesclous Sorter (⨀). In the preprocessing step, band-pass and CAR filters have been applied to raw data using the Tridesclous toolbox. During the spike sorting step, spikes were detected based on the threshold of 4 standard deviations (SD) of the background noise. All the multi-unit extraction was done automatically. So, the output of each recording session (*X*) from the spike sorting section would be *N* multi-units. Each multi-unit has *N*_1_ number of spikes. The details of the algorithm are illustrated in Appendix 1.

Further, to remove artifacts such as high frequency noise present in some channels, we have also considered two thresholds on the number of spikes from each multi-unit: the number of spikes in each multi-unit should be higher than 1600 and lower than 360000 for 900 seconds of recordings. After considering this criterion, the number of samples was reduced to 17 and TG3_D3 was removed. The number of multi-units extracted from each recording session is reported in Table. 1. In total, 53 multi-units (19 from WT, 34 from TG) were obtained.

**Table. 1.** Number of units (spike-trains) extracted after the spike sorting performed on the recorded data. There were 18 recording sessions (samples) from 8 rats (7 samples of WT rats, 11 samples of TG rats).

| Number | Subject code | Day | Total Recording Duration (Sec) | Number of units |
| --- | --- | --- | --- | --- |
| 1 | TG1 | D1 | 1624 | 4 |
| 2 | TG2 | D1 | 1488 | 8 |
| 3 | TG3 | D1 | 1821 | 2 |
| 4 | TG4 | D1 | 1563 | 2 |
| 5 | TG5 | D1 | 1140 | 1 |
| 6 | TG2 | D2 | 1720 | 7 |
| 7 | TG3 | D2 | 1801 | 2 |
| 8 | TG4 | D2 | 1668 | 3 |
| 9 | TG5 | D2 | 1592 | 2 |
| 10 | TG2 | D3 | 1801 | 3 |
| 11 | TG3 | D3 | 1610 | 2 |
| 12 | WT4 | D1 | 1049 | 2 |
| 13 | WT5 | D1 | 1506 | 2 |
| 14 | WT6 | D1 | 1417 | 3 |
| 15 | WT4 | D2 | 1590 | 2 |
| 16 | WT5 | D2 | 1528 | 3 |
| 17 | WT6 | D2 | 1575 | 5 |
| 18 | WT4 | D3 | 1416 | 2 |

#### Calculation of metrics reflecting Temporal Coding

In neuroscience, analyzing neural coding realizes the relationship between stimulus and neural response produced by the neuronal network. In temporal coding, the spike times of an individual neuron are considered to carry essential information. Here, to take into consideration the temporal complexity of spike sequences, we extracted different synchronization and distance metrics. These measures can compare two spike patterns and provide information about their relationship (similarity or dissimilarity). We have used three spike distance metrics to compute the spike train relationships among the units of each genotype group: 1-Van Rossum distance (VR-d), 2-Inter-Spike Interval distance (ISI-d), and 3-Event-Synchronization Distance (ES-d). VR-d is a metric for the dissimilarity of spike trains and measures the distance between two sets of spike trains by transforming them into continuous functions by convolving them with an exponential kernel^57^. ISI-d assesses the dissimilarity between spike trains based on instantaneous rate synchrony calculated by the time intervals between spikes. ES-d also measures spike synchrony albeit using a different approach. In contrast to ISI-d, it measures similarity instead of dissimilarity^58^. To calculate these metrics, isolated multi-units from each group of TG and WT have been concatenated to make a unified list of spike trains. The total number of spike trains used for each group is defined by *N*_*TG*_, *N*_*WT*_. In summary, all three spike distance metrics were calculated for each pair of spike trains. For more information on the mathematical description of each metric we refer to Appendix 2.

#### Machine Learning Classifier

Currently, the DL and ML models have a great potential to be used in hospitals and health care systems for clinical decision support in a variety of diseases including cancers^59,60^ and AD (image analysis)^61,62^. The number of dimensions in the feature space is one of the important parameters in an ML model that should be controlled to achieve reasonable accuracy while avoiding overfitting. In the final step of our model, we use unsupervised dimension reduction methods to reduce the feature dimensions. For this purpose, we applied PCA and the components carrying the highest explained varianSce were chosen. We have employed this approach for different numbers of components (1-9), and the best performance was achieved by considering three components (P=3). The training and validation steps were the same as the method discussed in the spectral analysis section.

#### Statistics

Mean ± SEM is provided for comparing different frequency bands. Statistical testing for frequency band power, PAC, and spike-train metrics, assumed non-parametric distributions and thus Mann Whitney U tests were used to evaluate differences between experimental groups.

## Results

### Low-Frequency Signal (LFS) Analysis

Several studies have shown that gamma oscillations are commonly disrupted in AD^39–41^. More specifically, fast gamma oscillations were shown to be diminished in the hippocampal CA1 region of a knock-in mouse AD model as well as the TgF344-AD rat model^39,40,63–65^. To evaluate if gamma and potentially other rhythms are impaired at an early stage in the hippocampus of TgF344-AD rats relative to WT controls, we computed the average PSD for different rhythms and specifically delta (1-4 Hz), theta (4-12 Hz), slow gamma (30-50 Hz) and fast gamma (50-100 Hz) (see Materials and Methods). In Figure 3, the average PSD for each pair of these bands is illustrated per recording session (WT: blue circles, TG: red circles). In addition, the diagonal panels show a univariate distribution that represents the marginal distribution of the data in each band (Fig.3). This illustration allows the comparison across the TG and WT groups for each pair of the four frequency bands to identify the ones that can reasonably discriminate between them. The results showed no apparent difference among TG and WT rats when the delta and theta frequency bands were considered with the samples of the two groups close to each other and a high degree of overlap between the distributions. On the contrary, when the slow and fast gamma oscillations were considered, it was immediately apparent that the separation between the distributions of PSD values across the two groups was substantial, promising higher discriminability in feature space and a better classification performance.

To better understand in which frequency bands differences occurred between the two groups, we plotted the power spectra, time-frequency plots, and normalized-power bar plots along with statistical analysis (Fig. 4). We found no significant differences in the average normalized power for delta and theta oscillations that were very similar across both groups (Delta; WT: 1.29 ± 0.03; TG: 1.09 ± 0.08; *p* > 0.01, Fig. 4 Theta; WT: 1.39 ± 0.19; TG: 1.2 ± 1.2; *p* > 0.01, Mann-Whitney U test, n = 7 WT and n = 11 TG recording sessions). Inversely, for both slow (30–50 Hz) and fast (50–100 Hz) gamma oscillations a significant reduction in average PSD observed (Slow gamma; WT: 0.13± 0.008; TG: 0.075 ± 0.008; *p* > 0.01, Fig. 4 Fast gamma; WT: 0.048± 0.0024; TG: 0.023± 0.0026; *p* > 0.01, Mann-Whitney U test, n = 7 WT and n = 11 TG recording sessions).

**Fig. 4.**
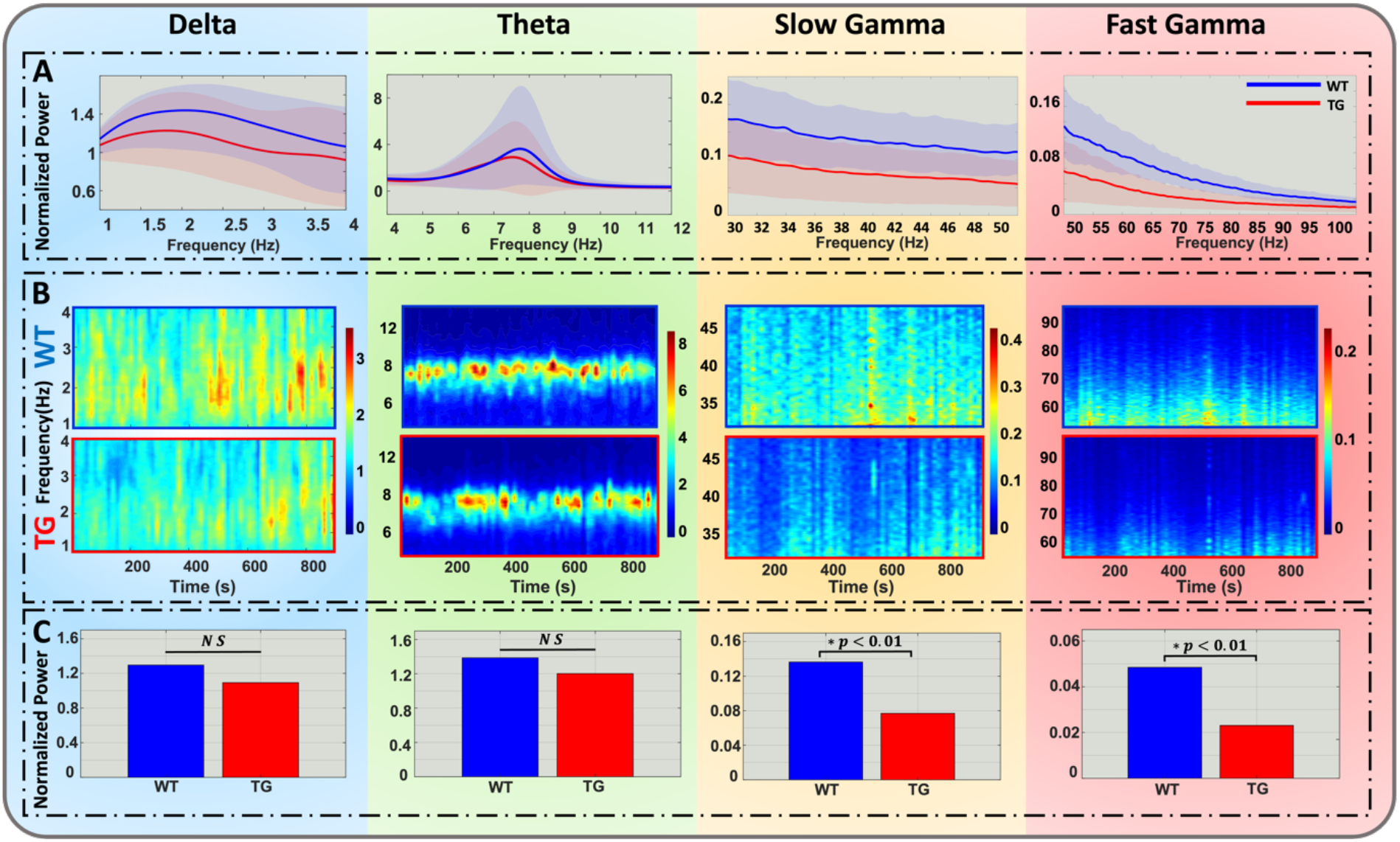
Comparison of the power spectral density (PSD) (A), time-frequency plots (B), and average PSD of 4 different frequency bands (Delta:1-4 Hz; Theta: 4-12 Hz; Slow Gamma: 30-50 Hz; Fast Gamma: 50-100 Hz) (C). (A) From left to right average mean PSD of the delta, theta, slow gamma, and fast gamma in WT rats (n = 7 recording session) and TG (n = 11 recording session). (B) The time-frequency spectra for for WT (upper panels) and TG rats (lower panels) for different frequencies of interest. (C) Normalized power for WT and TG rat in the frequency bands of interest. WT = wildtype, TG = TgF344-AD

Previous studies used electrophysiology recording in the same rat model have indicated an attenuation in the theta phase – gamma amplitude coupling (PAC) in the hippocampus of 8-9 months old animals^65,66^. To investigate if PAC, which plays an essential role in memory formation^29^, shows attenuation already at an earlier time point during disease progression, we assessed PAC across the WT and AD groups at 6 months of age. To this end, we calculated the modulation index (MI) capable of detecting cross-frequency coupling between two different frequency ranges of interest in a single signal as previously described by Tort et al.^32^. Example results from two TG and two WT rats are shown in Figure 5. The PAC spectrograms (Fig. 5B) revealed that large amplitude slow and fast gamma oscillations occur in the peak to falling phase of theta oscillations in WT; however, slow gamma oscillations seem to be attenuated in TG. Therefore, we have decided to compute the MI for fast and slow gamma among all samples (Fig. 5C). The analysis revealed no significant MI difference for fast gamma oscillation (Fig. 5C fast gamma; WT: 0.000342 ± 2.8 10^−5^; TG: 0.00037 ± 7 10^−5^; Mann-Whitney U test, n = 7 WT and n = 11 TG recording sessions); however, slow gamma coupling was significantly attenuated in TG (Fig. 5C slow gamma; WT: 0.00026 ± 4 10^−5^; TG: 0.000155 ± 1.7 10^−5^; Mann-Whitney U test p > 0.01, n = 7 WT and n = 11 TG recording sessions).

**Fig. 5.**
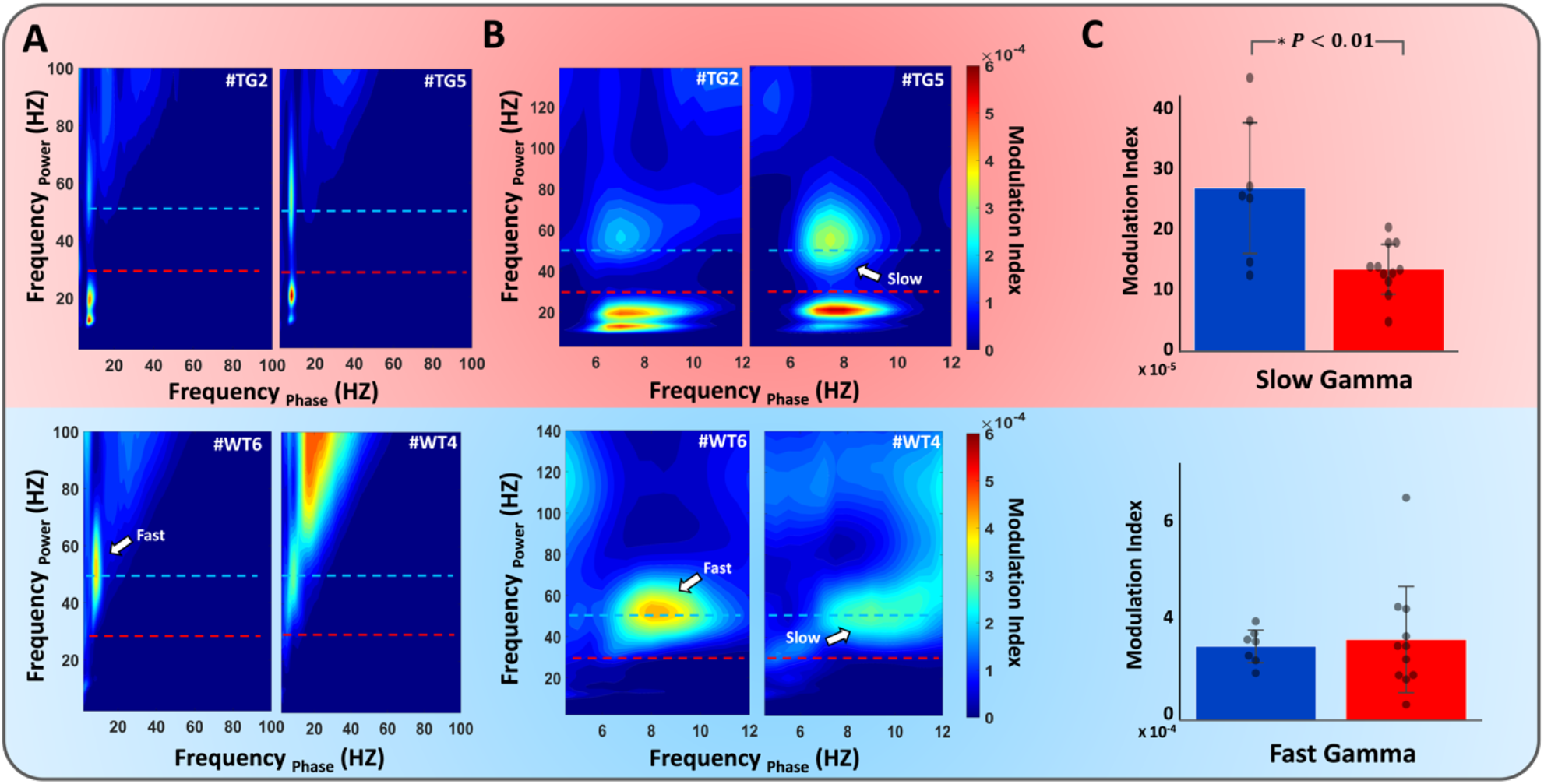
Slow gamma coupling was diminished in the hippocampal CA1 of 6 months old TgF344-AD rats. spectrograms of phase-amplitude coupling in the hippocampus were represented for two TG and WT rats by computing the modulation index to measure the coupling between desired frequency ranges. **A)** Comodulograms showing the phase and power of 0-100Hz frequency for two representative TgF344-AD (TG) rats (top row) two representative wildtype (WT) rats (bottom row). Modulation index values were coded by color and plotted according to theta and gamma frequency. The dotted lines denote 30 and 50 Hz borders between theta, slow gamma, and fast gamma oscillations. **B)** Comodulogram showing the phase of the theta band (5-12 Hz) in regard to the power in the 0-140Hz frequency band **C)** Modulation Index was computed for coupling of slow (30-50 Hz) and fast gamma (50-100 Hz) oscillations with theta rhythms on all recording sessions (WT: n = 7, TG: n = 11). Each black point represents each recording session of both groups. TG rats exhibited a significant decrease in slow gamma modulation index compared to WT (∗ *p* < 0.01).

Synchronous oscillations in various frequency ranges have been recorded in several nuclei of the basal BG and are thought to be an information processing mechanism. HVSs are 5–13 Hz spike-and-wave oscillations, which are commonly recorded in rats, and which have been reported in some recent studies where their occurrence in the BG has been investigated^67^. This distinct oscillatory pattern was present in the cortical area of TgF344-AD in previous studies and very rare in WT rats^68,69^. However, their occurrence rate was not investigated in the hippocampus of the AD rat model in the earlier stage of AD. HVSs are commonly characterized by an increase in their instantaneous power spectral density values; this must be incorporated into any HVS detection algorithm. The continuous wavelet transforms (CWT), and discrete wavelet transform (DWT) has been effectively applied for the detection of HVS-like nonstationary features such as sleep spindles and spike-wave discharges. We have investigated the alternations of HVSs in both time and frequency domains. The average time-frequency analysis of the two groups indicates no significant difference (Fig. 6A). We have further our analysis to make statistical inferences by calculating the HVS incidence metric (number of events/duration). Our results indicate there was no significant difference between the two groups in the occurrence rate of HVSs (Fig. 6B; WT: 2.7 ± 0.8; TG: 2.5 ± 0.6; Mann-Whitney U test, n = 7 WT and n = 11 TG recording sessions).

**Fig. 6.**
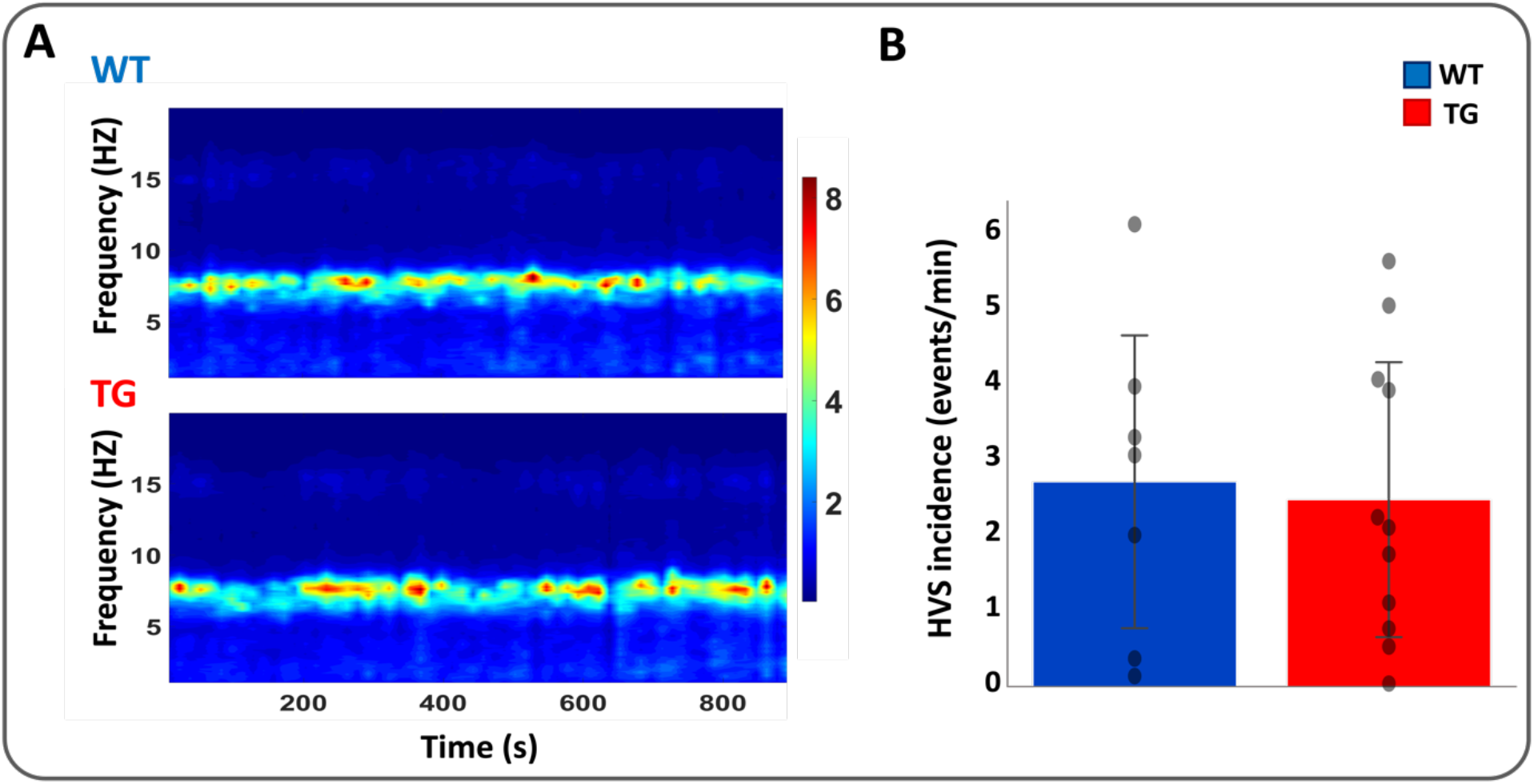
High-voltage spindles (HVSs) in the cortex of aged wild-type (WT) and TgF344-AD rats. **A)** Average time-frequency plots with a duration of 15 min of the hippocampus from WT and TG rats (WT: n = 7, TG: n = 11). Large negative deflections in the LFP and prominent bouts of high-power activity between 6 and 9 Hz in the spectrogram are HVS events. **B)** Comparison of HVS incidence rate between TG and WT rat models.

Subsequently, we proceeded to use ML classifiers (see Materials and Methods) to estimate their performance and ability to identify the rats from the WT and TG groups. To investigate how each frequency band affects the model’s accuracy, we first trained the ML classifiers with all frequency bands included. In Fig. 7A, the total accuracy among all samples is shown while Fig. 7B-C presents the performance separately in each group. To estimate the importance of each band, the fast gamma, slow gamma, theta, and delta oscillations have been removed one by one and their effects on the classification accuracy were calculated (Fig. 7D). When all frequency bands were used as input of the classifiers (RF, k-NN and SVM) the model performance was as expected maximum. However, the discrimination performance dropped significantly after eliminating fast and slow gamma oscillations indicating their importance in the classification of WT and AD rats.

**Fig. 7.**
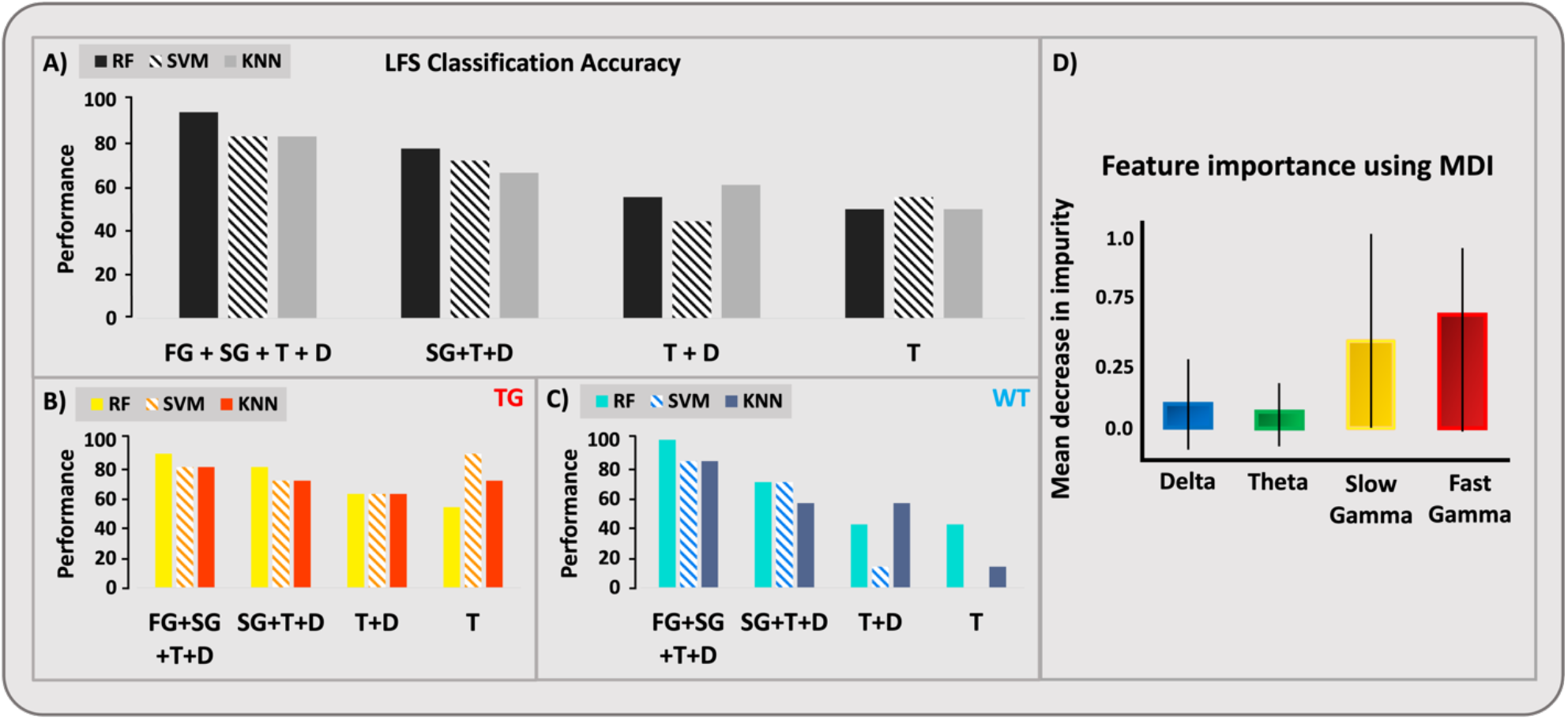
Classification accuracy of LFS model with three ML classifiers: k-nearest neighbors (k-NN), Support vector machine (SVM), and Random Forest (RF). Frequency bands: Fast Gamma (FG), Slow Gamma (SG), Theta (T), and Delta (D). (A) the overall accuracy. (B) the classification accuracy only for TG rats. (C) the classification accuracy only for WT rats. From left to right in each section, the frequency bands are removed from the classifier’s input one by one to see their effects on the accuracy. (D) the results of feature importance analysis. The MDI for each frequency band is calculated which shows the importance of slow and fast gamma oscillations.

Given that the number of samples from each rat was slightly different, the above-mentioned classification results could be susceptible to bias against rats with a low number of samples. Therefore, as a control, the results of the LFS classification task were reproduced with a selection of an unbiased sample (one sample from each subject) to ensure that the same results could be achieved. These results are available in the supplementary information (Fig S1, S2, S3).

### High-Frequency Signal (HFS) Analysis

Signals such as the spectral power in different frequency bands and phase-amplitude coupling, which we used in the LFS analysis, provide straightforward, measurable quantities that are also associated with specific cognitive mechanisms that are relatively well understood. In the case of HFS, the average firing rate of neurons is one of the quantities that is similarly straightforward to calculate and interpret. However, although the firing rate is a quantity that has been extensively used in neuroscience, its use is in most cases coupled with a stimulus (or task) and it is usually the stimulus response function that provides information about the underlying mechanisms and/or aberrations due to pathology. On the other hand, in the case of activity during relatively free behavior such as the one the rats in our study performed, it is very difficult to draw conclusions based on the firing rate alone. The huge variability in the base firing rates, which depend on cell types, brain areas or specific layers within them, as well as variability caused by the behavior itself, make the evaluation of spike trains based on the firing rate problematic and difficult to understand. To this end, spike train distance metrics have been developed that provide a powerful alternative approach to evaluate spiking activity in more general conditions. Moreover, depending on the spike-trains distance metric used, we can deduce information whether changes depend on underlying principles of neural coding such as rate or temporal codes. Here, we used three different spike-train distance metrics: VR-d, a parametric distance based on the temporal structure of a pair of spike trains, ISI-d, a parameter-free spike distance metric that extracts information from inter-spike intervals by evaluating the ratio of instantaneous firing rates, and ES-d, a complementary measure that is sensitive to spike coincidences. Given that these metrics depend on different characteristics extracted from the spike trains, the performance of each metric in the classification task can provide information about the underlying mechanisms being disrupted in AD.

In Fig. 8 A and B, the spike distances among all spike patterns calculated in the TG and WT groups are shown. It is immediately obvious that the patterns for each metric show differences and similarities across the two groups. VR-d is a parametric spike train distance which is based on the temporal structure of two pairs of spike trains. In Fig 8 (left panels), we can observe that the VR distance between WT multi-units is generally small. However, the distance between TG multi-units is not consistent, and some spike trains have considerable distance from the others. ISI-d is a parameter-free spike distance metric that extracts information from inter-spike intervals by evaluating the ratio of instantaneous firing rates. In Fig. 8 (middle panels), the ISI distances among both TG and WT spike trains demonstrate similar values and variability and thus it is not immediately obvious if this metric can provide information for classification. A major weakness of ISI-d is the fact that it is not well suited to track synchrony changes based on spike coincidences. On the contrary, ES-d is a complementary metric that is sensitive to spike coincidences while sharing the fundamental advantages of ISI-d. In Fig 8 (right panels), like ISI-d, the values and variability across the two groups are similar and thus it is difficult to judge any difference between the two groups.

**Fig. 8.**
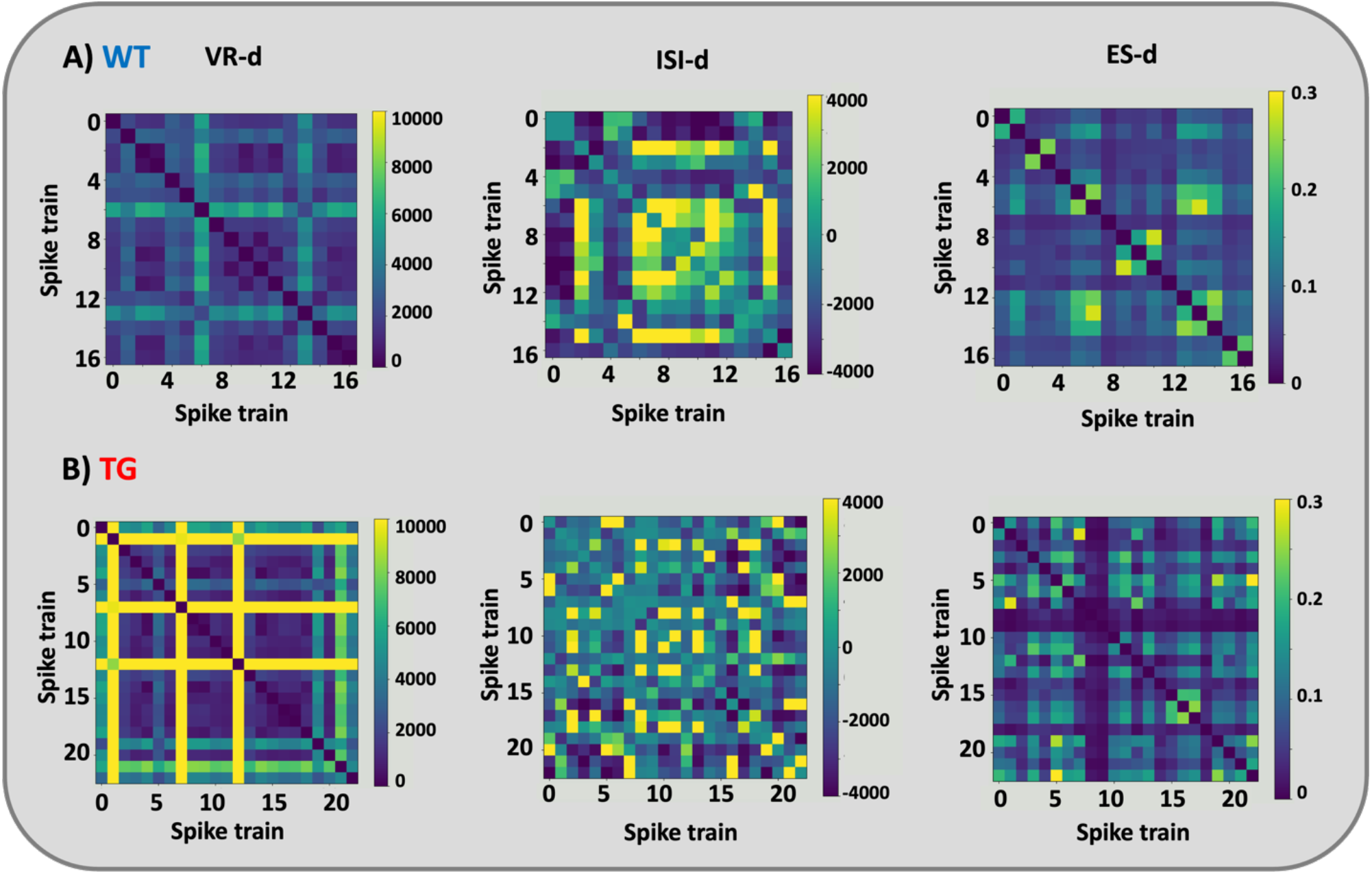
The spike distance metrics for individual spike responses in (A) TG and (B) WT groups. From left to right, van Rossum distance (VR-d), inter-spike interval distance (ISI-d) and event synchronization distance (ES-d). The blue color shows more similarity, and the yellow color indicates how far the two spike trains are.

As indicated in the Materials and Methods, dimensionality reduction via PCA was subsequently performed and the three components explaining the highest variability were used as features for further analysis. In Figure 9, the results of the PCA are presented in 3D feature space (Fig. 9). Using this representation, it becomes easier to observe that in the case of VR-d the data points of the WT group are very close to each other forming a cluster that is separated from the datapoints of the AD group. Thus, VR-d is a promising metric for classification. For the other two metrics, ISI-d and ES-d, the data points of the two groups are intermixed and thus it is more difficult to predict the results of classification.

**Fig. 9.**
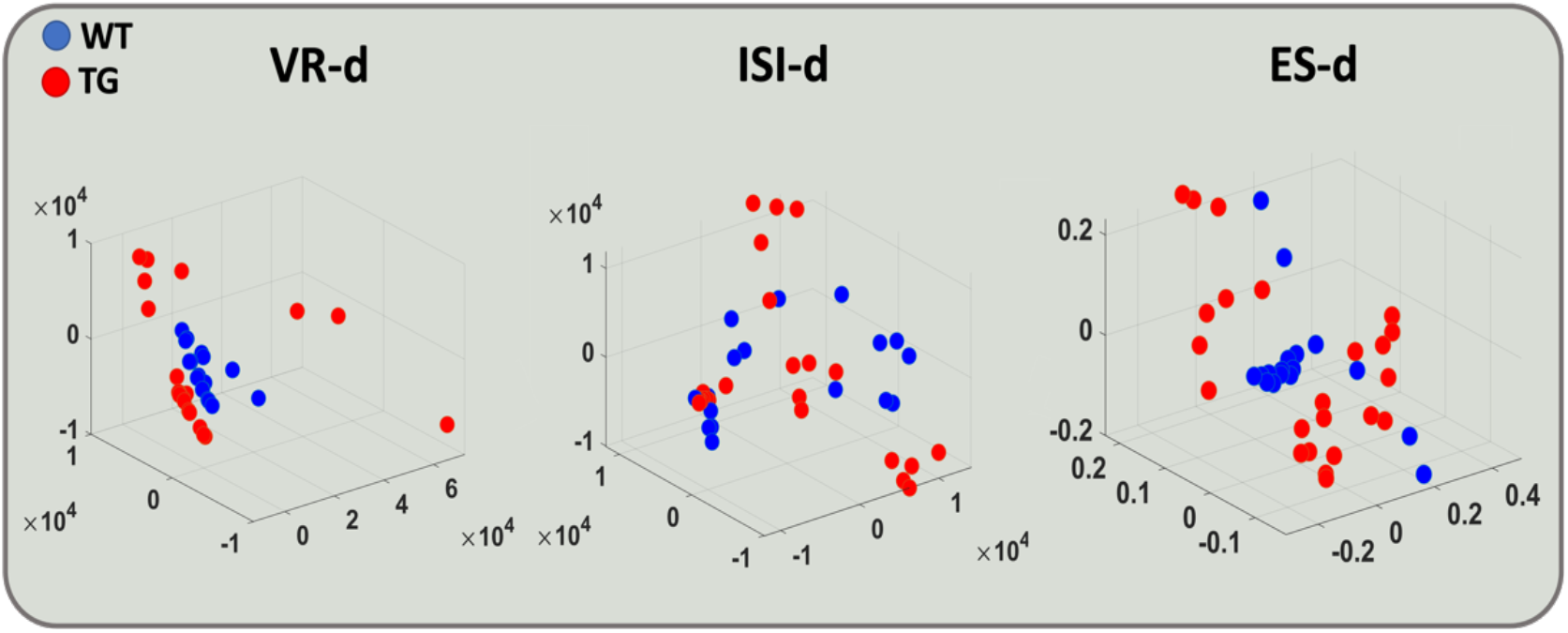
The feature space of each distance metric in 3D space. The red and blue dots represent TG and WT rats respectively. Each figure is the result of the application of PCA algorithm on the matrix of spike train distances (Fig. 7). Three prinicipal components from each spike distance matrix were considered as the inputs of the classifiers.

Using the features extracted from PCA analysis we then proceeded to using classifiers and namely SVM, k-NN, and RF to classify the two groups of rats based on HFS (Fig. 10). Note that for k-NN, different values of *k* from 3 to 13 have been tested to evaluate the model. However, we have only reported the result of *k* = 5 as the best result. The classification results were compatible with the observed separation of groups in the feature space (Fig. 9) with the features based on the VR-d providing the highest total classification accuracy for all classifiers while the ISI-d had the lowest total performance (Fig. 10A). Notably, when the performance for each group was plotted separately (Fig. 10 B-C) a slightly different behavior across groups was observed. For the WT group VR-d showed clearly the highest accuracy reaching 100% for SVM and k-NN classifiers while ISI-d performed very poorly; ES-d had intermediate performance. For the TG group, on the other hand, the maximum classification accuracy was in general less but comparable for each metric.

**Fig. 10.**
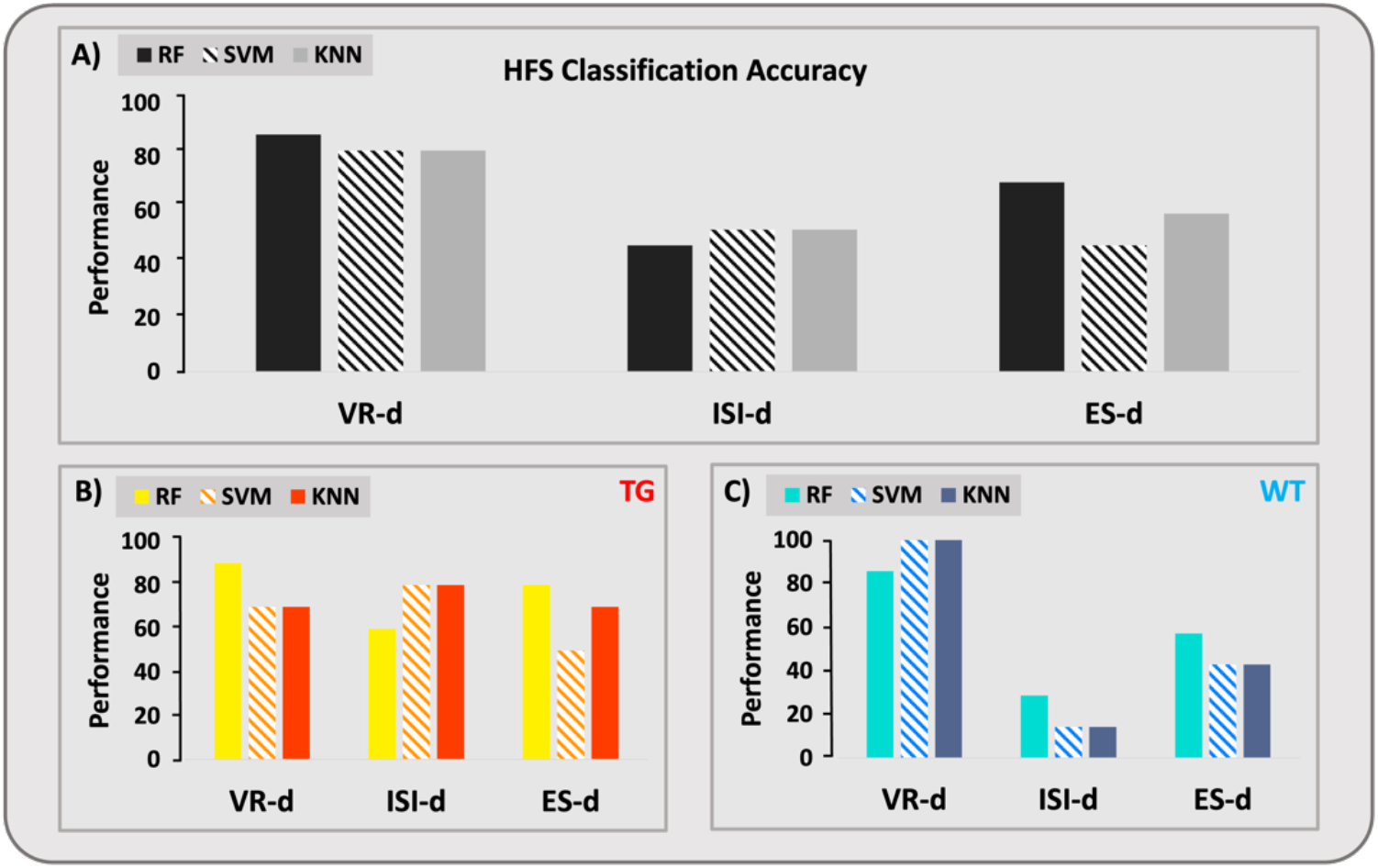
Classification accuracy for three different classifiers: RF, SVM, k-NN. LOSO-CV was used for training and testing individual classifiers. (A) the total performance for the classification of TG and WT samples is presented. The classification performance for (B) TG rats and (C) for WT rats separately.

### Concatenation of HFS and LFS Analysis

As we reported in the previous sections, we have used two types of analysis to extract features from low and high-frequency signals respectively. Because each analysis extracts features with distinct characteristics, the LFS and HFS models do not share information. Therefore, it is possible to combine LFS and HFS information by stacking these models. The simplest form of stacking can be described as an ensemble learning technique where the predictions of multiple classifiers (referred as level-one classifiers) are used as new features to train a meta-classifier^70^. Fig. 11A shows how LFS and HSF models have been stacked as level-one classifiers and logistic regression as meta-classifier. The predictions of level-one models (*P*_1_, *P*_2_) are stacked and used as input to train the final prediction’s meta-classifier. To stack models, the best model of HFS (RF model using VR-d features) and the best model of LFS (RF model) were chosen. The meta-model is often a simple model, providing a smooth interpretation of the predictions made by the base models. As a result, linear models are often used as the meta-model, such as linear regression for regression tasks (predicting a numeric value) and logistic regression for classification tasks (predicting a class label). Although this is common, it is not required. Logistic regression is a statistical technique used to estimate the probability of a discrete outcome given an input variable. Also, it is best thought of as linear regression modified for classification problems^71^. The major distinction between linear and logistic regression is that the range of logistic regression is limited to values between 0 and 1. Table 1 provides the classification performance of the proposed ML method. According to the Classification results, we have achieved 80.23% and 94.44%, respectively, using high and low-frequency features. However, using the combination of high and low-frequency features using stacking models, the accuracy was 100%, and the feature space of two classes was easily separable with a single line (Fig. 11B).

**Fig. 11.**
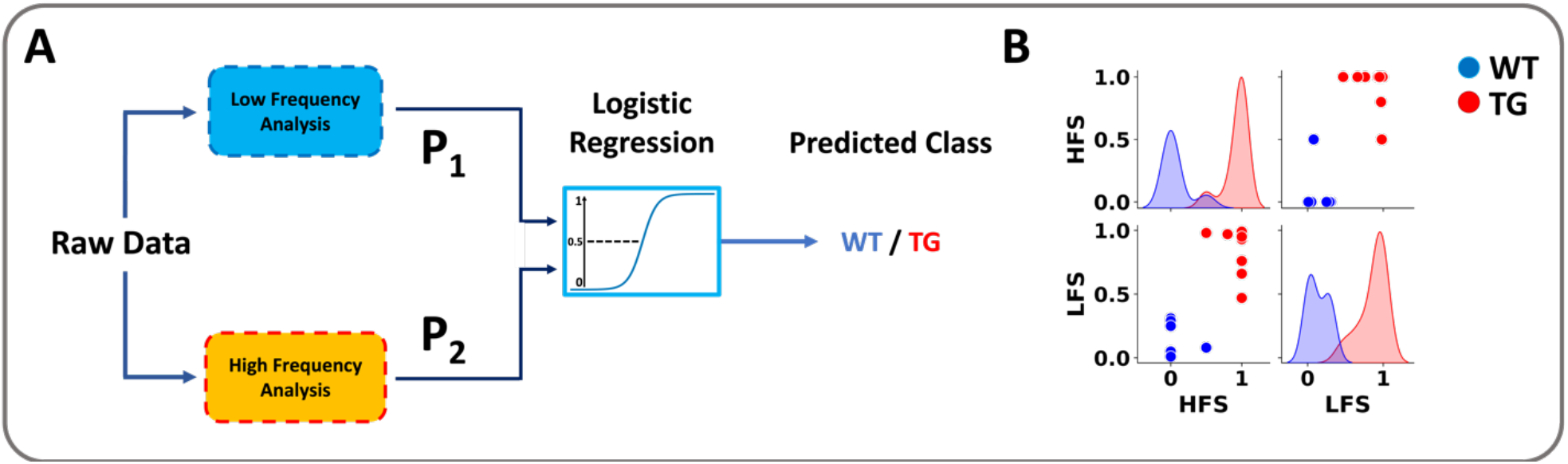
**A)** Stacking model pipeline is represented. *P*_1_ is the probabilistic prediction of the LFS model and *P*_2_ is the average prediction of each mouse’s spike train using the HFS model. Next, *P*_1_ and *P*_2_ are used as input of meta-model (logistic regression). **B)** The feature space of meta-model for classification. The red and blue dots represent TG and WT rats respectively.

**Table 1.** Comparison of the classification models

| Model | TG | WT | Total |
| --- | --- | --- | --- |
| LFS (RF) | %90.9 | %100 | %94.44 |
| HFS (RF - VR-d) | %90 | %85.71 | %80.23 |
| Stacked | %100 | %100 | %100 |

## Discussion

Alzheimer’s disease is characterized by the progressive accumulation of amyloid-β plagues and neurofibrillary tau tangles in the brain and is gradually leading to cognitive decline through multiple stages starting with mild memory loss, increasing deficits in daily life activities, loss of executive functions, and eventually dementia^72,73^. To date, no therapies exist able to stop or decelerate this devastating condition, while diagnosis is possible only years after the onset of pathophysiological events, presenting a major challenge towards developing successful therapeutic approaches. Thus, current efforts are focusing in identifying early disease biomarkers able to provide prognostic features that will allow patient identification at early time points.

In this study, we recorded electrophysiological signals (LFPs and neuronal spiking activity) from the hippocampus of freely moving TgF344-AD rats and WT littermates, at an early phase of disease progression. Previous studies that characterized in detail the pathophysiology of this animal model, have indicated its great potential over other murine models, because it gradually expresses all hallmarks of AD in humans including the rarely observed (in murine models) spontaneous accumulation of endogenous hyperphosphorylated tau (pTau) and neurodegeneration^74,75^. Very importantly, the relatively slow disease progression dynamics in this model that simulate different stages of AD in humans, offer remarkable opportunities for identifying novel AD biomarkers, understanding the underlying mechanisms of pathology and cognitive decline, and testing new therapeutic approaches. At the period we performed our recordings (6 months old rats), only mild Aβ accumulation is observed in the cortex and hippocampus, cognitive deficits are by and large absent, and thus this time-point is accepted to simulate the early pre-symptomatic phase of AD in humans. We hypothesized, that subtle changes in electrophysiological activity in the hippocampus, could be potentially used to classify individual rats across the two genotypes, AD and WT.

The hippocampus, an area amongst the first to be affected in Alzheimer’s patients and strongly implicated in learning and memory processes, has been consistently shown to exhibit abnormal morphology and function in AD patients, with a remarkable reduction in volume as well as aberrant activation during cognitive tasks^68,69,76,77^. Consistent with these clinical observations, pre-clinical and translational research in transgenic animal models of AD pathology have repeatedly shown hippocampal dysfunctions and in particular abnormalities in oscillatory brain rhythms. Accumulating evidence suggests that disturbances in synaptic and neuronal function are leading to impairments of coordinated activity at early pre-symptomatic stages of the disease with major targets the brain networks that support memory and cognition^73,78^. Aberrations in the activity of these brain circuits range from subtle changes in oscillatory rhythms to more profound alterations that are reflected in EEG signals at later stages of the disease. Such impairments in brain rhythms, considered to underlie compromised hippocampal-dependent cognitive function, were also shown in pre-clinical animal models of AD before the onset of behavioral deficits^79–81^.

To identify potential early aberrations in electrophysiological signals in the hippocampus of our TgF344-AD rat model, we first performed analysis of the low frequency signals (LFPs). Two different frequency ranges of gamma oscillations thought to be generated in the hippocampus were analyzed: slow gamma (25–50 Hz) and fast gamma (50– 100 Hz)^24,82–84^. We found that both the slow as well as the fast gamma oscillations were reduced in the AD rats in comparison to WTs (Fig. 4). This is consistent with findings in which gamma oscillations were found to be disrupted in AD patients^38^, in mice^39,40^, but also in some previous experiments in the same transgenic rat model, albeit performed at a slightly later stage of disease progression (8-9 months of age)^65,66^. Noteworthy, those studies reported significant decreases in the slow gamma power, while in our study – performed at an earlier AD stage - we, in addition, observed strong decreases in the fast gamma oscillations. We speculate that this difference could potentially relate to compensation processes that get activated at these slightly later stages and are resulting in cognitive resilience that is persistent until much later time points^85,86^. On the other hand, our analysis of HVSs indicates no significant difference between the HVSs occurrence rate between two groups. Earlier studies have shown that HVSs would increase during the AD in the cortical region of TgF344-AD rats^68,69^. These spindles can reach the hippocampus by the neocortical-entorhinal cortex path^87^. Therefore, our assumption would be although these spindles have become widespread in the cortical region in the previous studies, they still have not reached the hippocampus in the earlier stages of AD.

In the hippocampus, gamma oscillations are nested within the slower theta oscillations with this co-modulation being proposed as a mechanism that recruits cell assemblies throughout the theta cycle^88^. The coupling between the phase of the theta with the amplitude of gamma oscillations has been demonstrated to support cognitive processes and memory formation in humans^31^, monkeys ^29^, and rodents^32,83,89^. To probe potential changes in these coupling in the TgF344-AD rat model we performed analysis of cross-frequency coupling (Fig. 5). We found a significant reduction of the modulation index in the theta phase with slow gamma but not the fast gamma amplitude in the AD rats. Low-frequency gamma is associated with gamma locking and signal transfer in the hippocampus from CA3 → CA1 (Fig. 2A)^24^. Previous research in TgF344-AD has found a reduced fidelity in the activity of place cells recorded from the hippocampus CA2/3 while place cells recorded from CA1 showed persistent fidelity until much later stages of the disease^90^. Thus, the reduced theta-phase slow gamma amplitude coupling we observe could stem from such early aberrations in the CA3 circuit. On the other hand, the absence of changes in theta-phase to fast gamma amplitude coupling, which relates to the signal transfer from MEC → CA1, indicates that this part of the circuit is able to function in a relatively normal fashion in the earlier stages of AD, although some synaptic deficits were found in vitro^91^.

Using the information and insights we obtained from the frequency band power analysis, we performed classification using combinations of the power in different frequency bands in our effort to understand the contribution of each band on classification accuracy. As expected from visualizing the pairwise combinations of the PSD values for different frequency bands (Fig. 3), our results indicated that the best accuracy is achieved when average power in the gamma band is one of the classifier’s inputs while if only theta and delta band information is used, classification is close to chance levels. Comparing the performance of different ML classifiers, we observed that their performance was similar on average, with the random forest achieving the highest total classification (Fig. 7). The importance of gamma band activity in classification across the groups is in line with previous findings^39,40,63–65,81,92^ suggesting that gamma oscillations are essential in spatial memory and get disrupted during early stages of AD. Moreover, this finding provides support to suggestions that gamma frequency entrainment or reactivation have positive effects and can be developed towards potential treatments for AD ^93,94^.

Having established that low frequency electrophysiology signals carry important information for classification, we then performed analysis of the high frequency signals by performing spike sorting to extract the spike trains of activity from different cells. Then, we calculated three spike-train distance metrics (VR-d, ISI-d, ES-d) across pairs of units in the TG and WT animals and used this as features for classification of the genotype. We found that VR-d provided the highest classification performance with over 80% total accuracy while the ISI-d and ES-d performance was on average close to chance levels. Interestingly, the ISI-d performance was dissociated between the two genotype groups showing high classification performance in TG animals but low in WT. In our experiments, we have recorded data from the hippocampus CA1 and CA3 regions ^95–97^ while the rats were freely moving in a linear track. We thus conjecture that aberrations in spike-trains driving the classification are related at least in part to place cell activity. Many recent studies have shown disruption of place cells activity in AD ^39,40,90^. A study by Galloway and colleagues, explicitly studied place cell activity in the hippocampus of TgF344-AD rats between 12 and 20 months of age ^90^. They found age related reductions in the fidelity of place cell firing in CA2/3 but not CA1. These subfields of hippocampus are part of the well-known trisynaptic loop which includes the dentate gyrus, CA3 and CA, with each of these subserving a different function^98^. The pyramidal neurons of CA3 exhibit burst activity and project to the CA1 pyramidal neurons via the Schaffer Collaterals. Noteworthy, the synapses of these CA3 neurons on CA1 dendrites are highly modifiable and very dynamic ^99^. In addition, approximately one third of synaptic connections in CA1 are received directly from the layer III of entorhinal cortex via the perforant pathway in contrast to CA2 and CA3 neurons, which receive direct projections from entorhinal cortex layer II ^100,101^. Given that previous studies have already shown that CA1 place cells are able to demonstrate normal place fields in the absence of CA3 input, albeit not without inputs from the layer III of the entorhinal cortex^102,103^, it would be perhaps possible to normalize hippocampal activity in AD by stimulating these layer III projections from the entorhinal cortex to CA1.

To further understand if the aberrations in hippocampal spiking activity are providing the same (or additional) information for classification as the low frequency signals, we decided to perform a concatenation of the HFS and LFS analyses into a combined ML model. We found that stacking information from the two analyses, boosted the classification to 100% accuracy. This could indicate that although a lot of information in these frequency ranges may be shared (e.g., due to the phase locking of spiking to the theta and/or gamma rhythms), additional independent information can help the accurate classification. Thus, our analysis is consistent with previous findings indicating that that connectivity between CA3 and CA1 subfields of the hippocampus is disrupted due to aberrations in CA3 activity while CA1 seems resilient.

In conclusion, the results of this paper indicate that with a combination of ML models, feature extraction methods based on similarities and distances in the neural responses (high-frequency components of the neuronal data) and time-frequency analysis of LFP signal (low-frequency components of the neuronal data) recorded from the rat’s hippocampus, we can classify healthy and AD rats at an early, pre-symptomatic stage of Alzheimer’s disease. These promising findings provide a clear indication that information from a combination of signals and electrophysiological measurements, could accelerate diagnosis of AD and assist scientists in the development of more effective therapies.

## Acknowledgments

This study was supported by the Fund for Scientific Research Flanders (FWO) (grant agreement G048917N to GAK) (G045420N to MV) and the Stichting Alzheimer Onderzoek (SAO-FRA-2018003 to MV).

## Author Contributions

M.v.B and G.A.K designed the electrophysiology experiments and G.A.K supervised the data acquisition. G.A.K. and M.A designed the simulations and M.A supervised the data analysis. F.M and M.A performed the high-frequency analysis. M.M, F.M, and M.A performed the low-frequency analysis. M.v.B performed the electrophysiology experiments and data acquisition in freely moving rats with the assistance of L.K. The initial draft of the manuscript was written by F.M, M.M, M.A, G.A.K. All authors edited the manuscript and accepted the final version.

## Appendix 1

The algorithm for spike sorting and multi-unit classification model

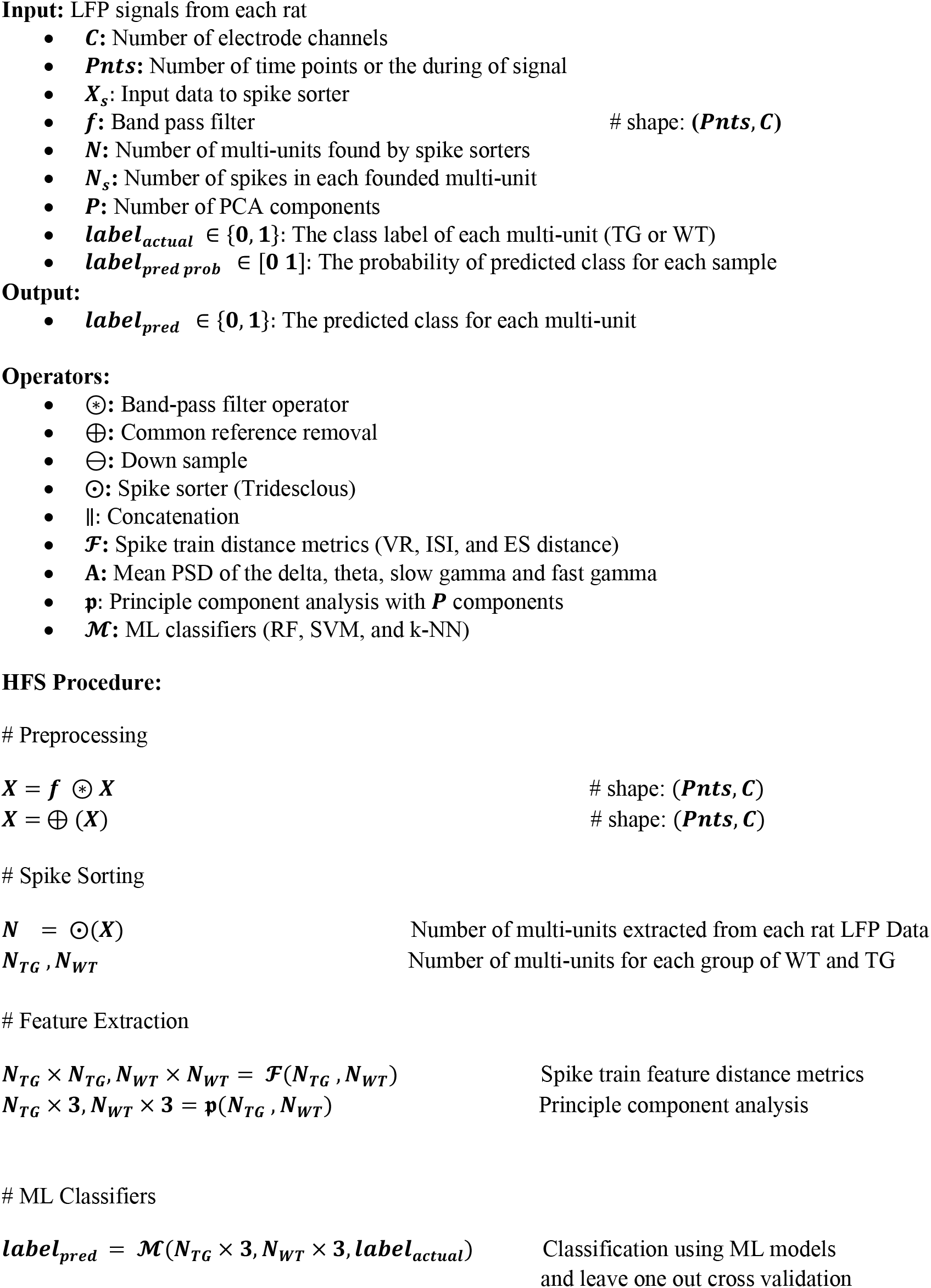
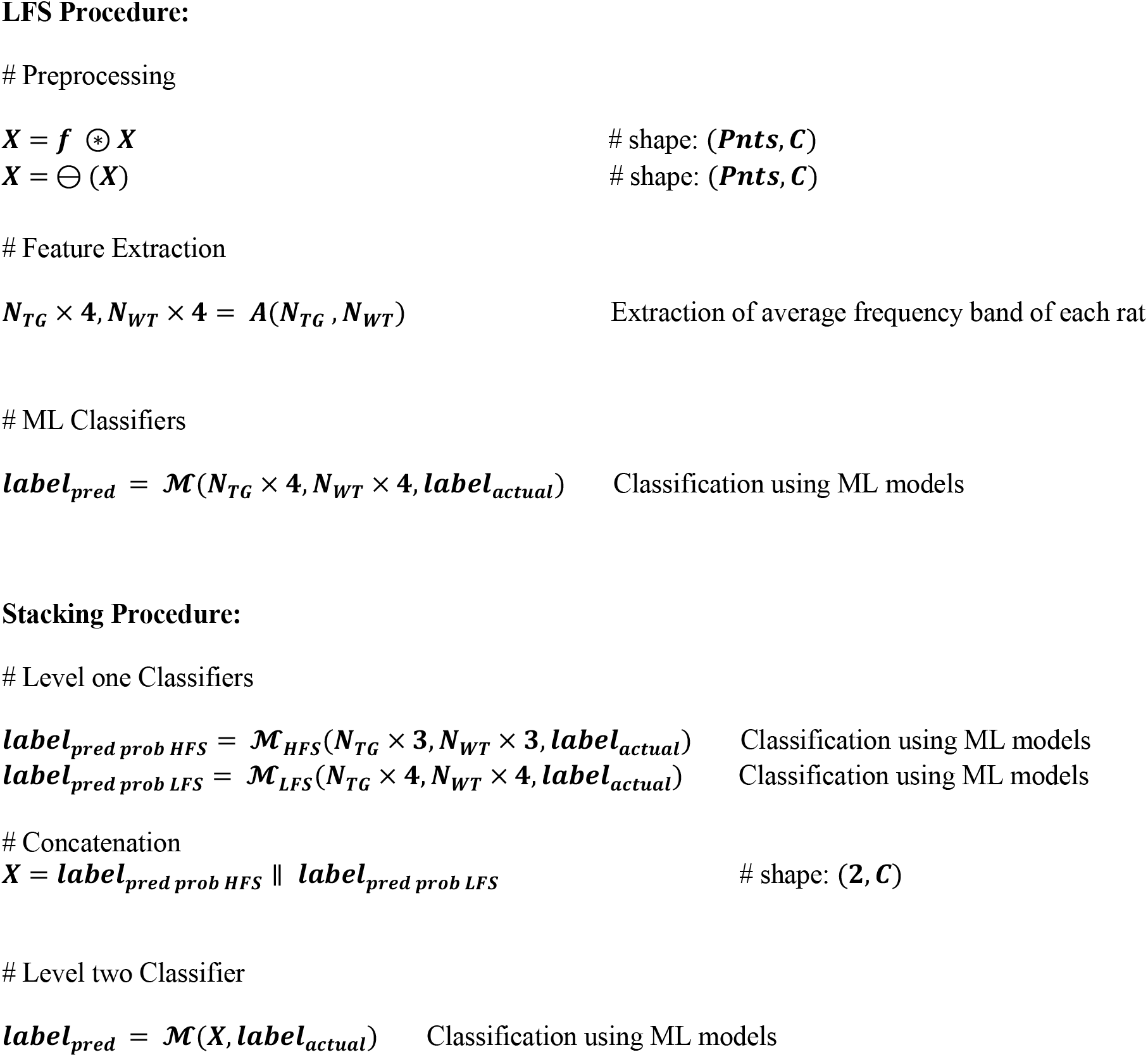

## Appendix 2

The mathematical description of individual metrics.

### Van Rossum distance (VRd)

The Van Rossum metric estimates the distance of two spike trains and has been proved to be a reliable metric as the distance between multiple responses to the same stimulus is usually smaller than the distance between responses to the other stimuli. The discrete spike trains x and y are transformed into continuous functions through convolving each spike *t*_*K*_ with an exponential kernel 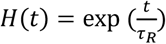. *τ*_*R*_ is the time constant and H is the Heaviside step function. After proper tradeoff, *τ*_*R*_ = 0.05 was selected. From the resulted waveforms *x*(*t*) and *y*(*t*), the VRd, *DVR* can be calculated as: [33]

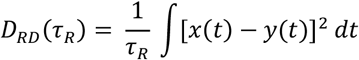

### Inter spike interval distance (ISI-distance)

The ISI-distance, *DI*, measures the immediate rate difference between spikes. It is based on a time-resolved profile that defines a dissimilarity value for each instance. We assign *t*, the time of the previous spike to each moment to obtain this feature:

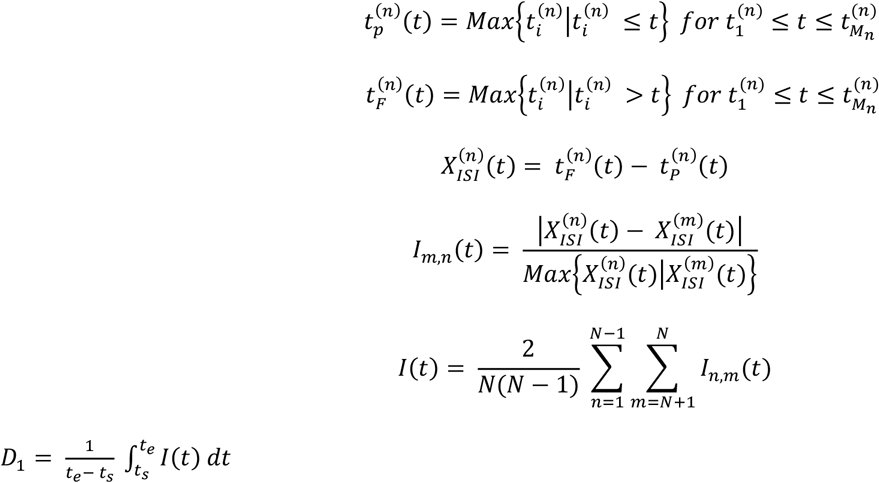

### The Event-Synchronization distance (ESd)

The Event-Synchronization distance is based on the event relative timing. In this article, the event is defined by a spike in the spike patterns. The spikes *ti* and *tj* in the separate spike trains are synchronous only if their time difference is less than the delay time *τij*, which is obtained by:

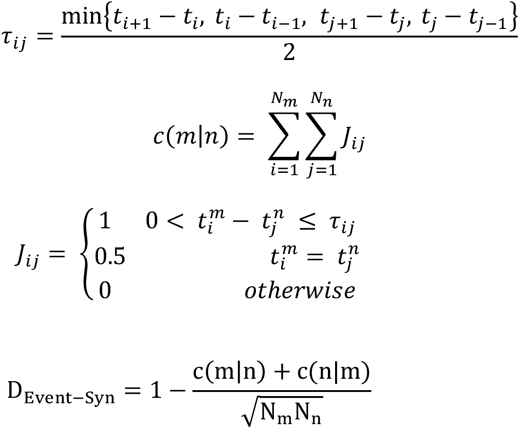

### Mean Decrease Impurity (MDI)

Gini importance measures the importance of variable X in predicting variable Y in the random forest for all nodes averaged over all *N*_*T*_ trees in the forest ^104^. This measure is also called Mean Decrease Impurity, here denoted by *IncNodePurity*:

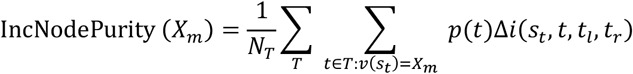

where *v*(*s*_*t*_) is the variable used in split *s*_*t*_ and Δ*i*(*s*_*t*_, *t, t*_*l*_, *t*_*r*_) is the impurity decrease of a binary split *s*_*t*_ dividing node *t* into a left node *t*_*l*_ and a right node *t*_*r*_:

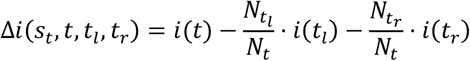

where *N* is the sample size, 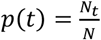the proportion of samples reaching t, and 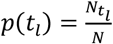and 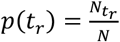are the proportion of samples reaching the left node *t*_*l*_ and the right node *t*_*r*_ respectively.

## Supplemental Information

**Figure S1.**
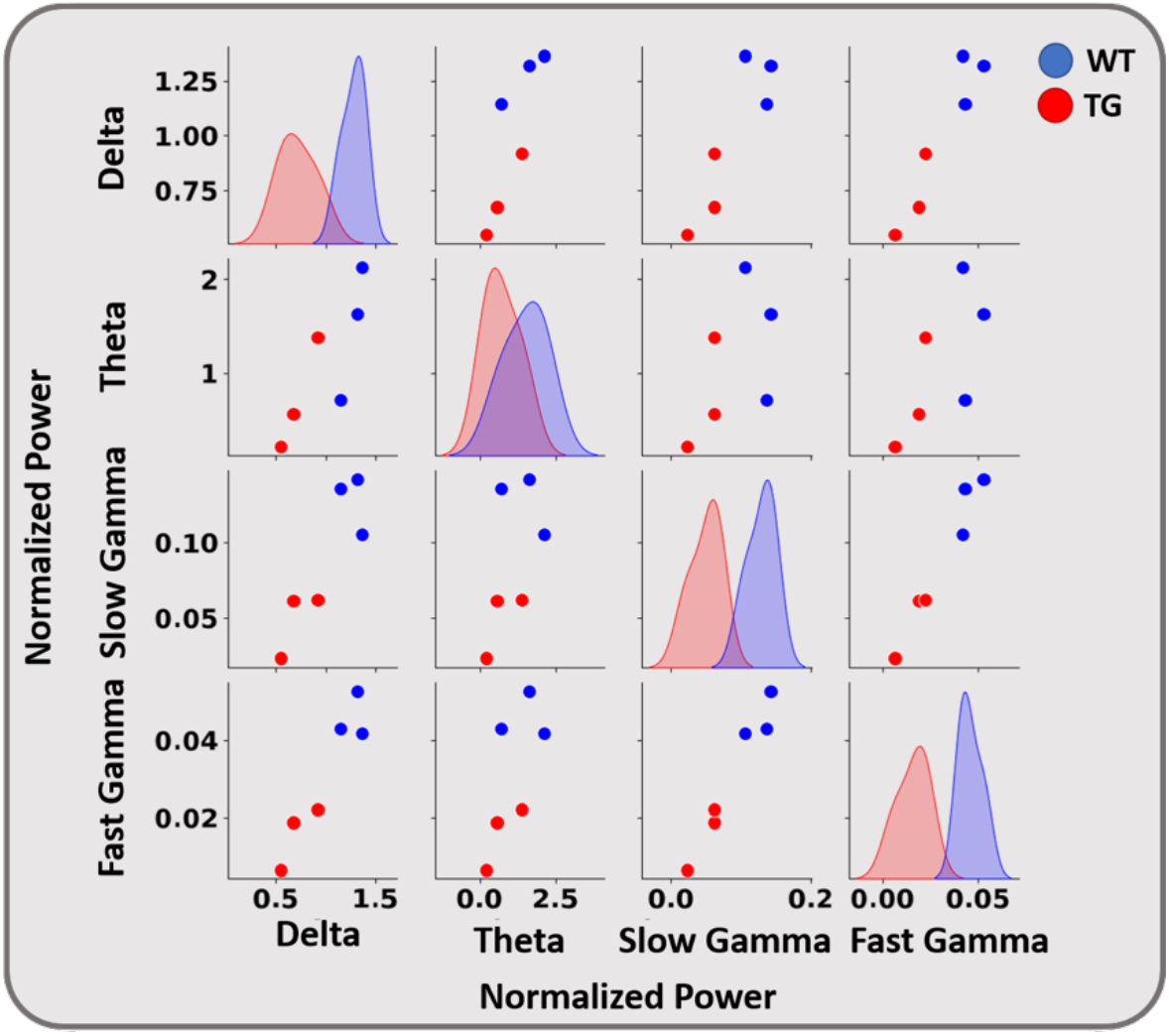
Pairwise comparison of the normalized power in delta (1-4 Hz), theta (4-12 Hz), slow gamma (30-50 Hz), and fast gamma (50-100 Hz) for TgF344-AD (TG) and Wildtype (WT) groups. Each point represents the average PSD of a single sample in the specified band. The diagonal plots are the univariate distributions that show the amount of overlap between the distribution of two groups in individual frequency bands. The other plots compare WT and TG for specified frequency bands. Blue/Red points are recording session (samples) of WT (n = 3) /TG (n = 3), respectively (TG1, TG2, TG3, WT4, WT5, WT6).

**Figure S2.**
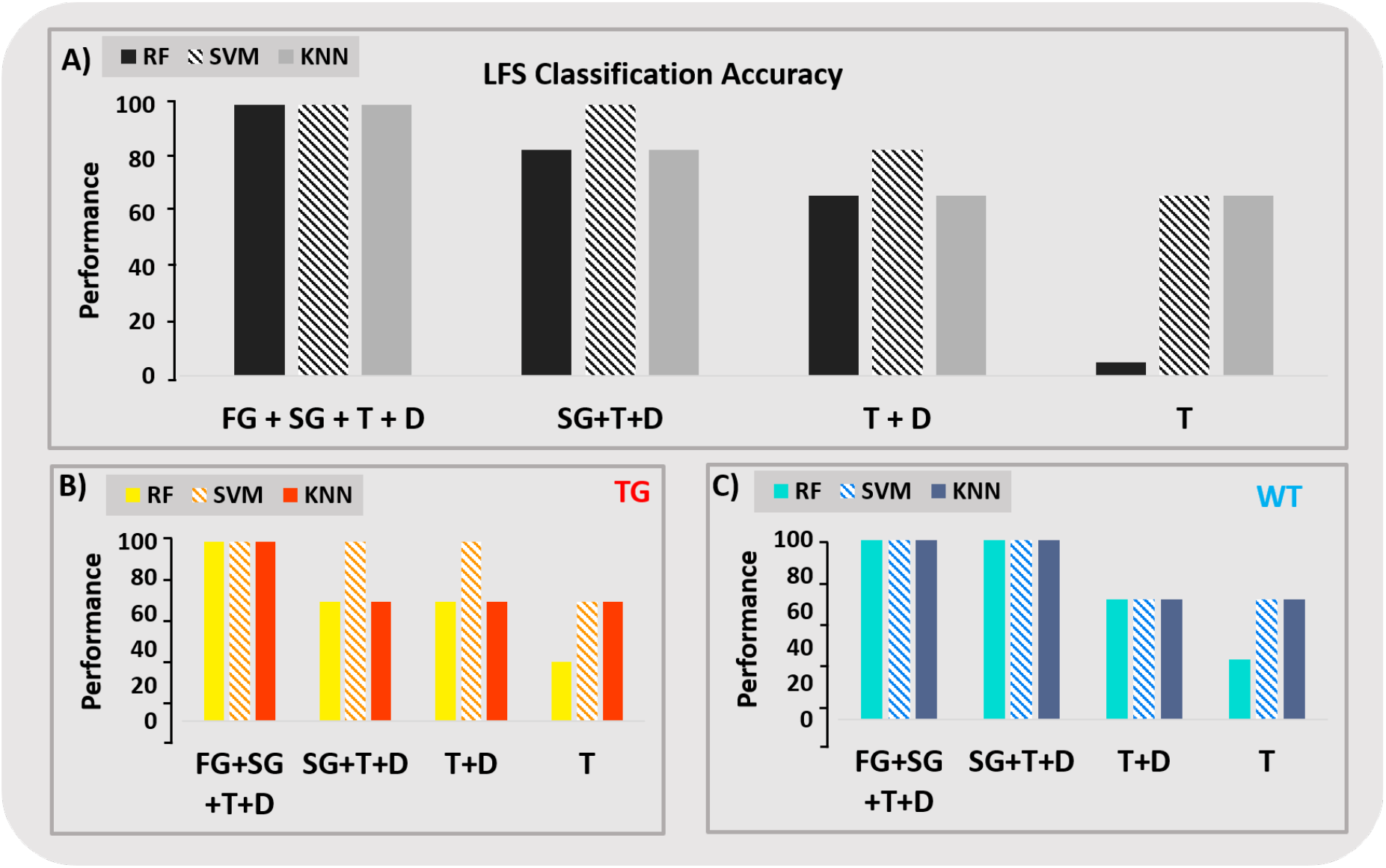
Classification accuracy of LFS model with three ML classifiers: k-nearest neighbors (k-NN), Support vector machine (SVM), and Random Forest (RF) for 6 rats from both groups of TgF344-AD and WT (TG1, TG2, TG3, WT4, WT5, WT6). Frequency bands: Fast Gamma (FG), Slow Gamma (SG), Theta (T), and Delta (D). (A) the overall accuracy. (B) the classification accuracy only for TG rats. (C) the classification accuracy only for WT rats. From left to right in each section, the frequency bands are removed from the classifier’s input one by one to see their effects on the accuracy.

**Figure S3.**
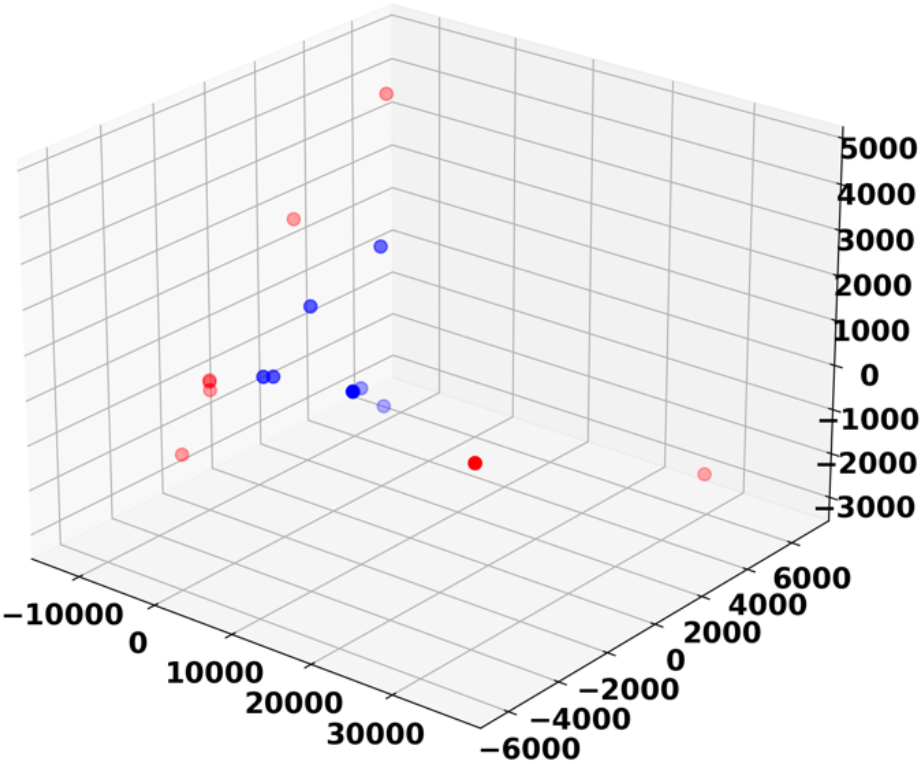
The feature space of VR-d by considering 6 rats from both groups of TgF344-AD and WT (TG1, TG2, TG3, WT4, WT5, WT6) in 3D space. The red and blue dots represent TG and WT rats respectively. Each figure is the result of the application of PCA algorithm on the matrix of spike train distances (Fig. 7). Three prinicipal components from each spike distance matrix were considered as the inputs of the classifiers.

